# A high-resolution haplotype pangenome uncovers somatic hybridization, recombination and intercontinental migration in oat crown rust

**DOI:** 10.1101/2024.03.27.583983

**Authors:** Eva C. Henningsen, David Lewis, Eric Nazareno, Yung-Fen Huang, Brian J. Steffenson, Brendan Boesen, Shahryar F. Kianian, Eric Stone, Peter N. Dodds, Jana Sperschneider, Melania Figueroa

**Affiliations:** Commonwealth Scientific and Industrial Research Organisation, ACT Australia; Australian National University, ACT Australia; University of Minnesota, Saint Paul, MN U.S.A.; National Taiwan University, Taipei, Taiwan; Cereal Disease Laboratory, USDA-ARS, St. Paul, MN U.S.A.

**Author notes:** Corresponding author: M. Figueroa.

**Keywords:** haplotype, mating type, pangenome, *Puccinia coronata*, somatic hybridization, virulence

## Abstract

Basidiomycetes like rust fungi have complex reproductive cycles and dikaryotic life stages which influence their population structure and evolution. *Puccinia coronata* f. sp. *avenae* (*Pca*), the causal agent of oat crown rust, is a pathogen of global economic importance. To investigate the genetic diversity of the species, as well as the role of mating type system and nuclear exchange (somatic hybridization) in host adaptation of *Pca* we acquired whole genome sequencing data of Taiwanese and Australian isolates, adding to existing data for USA and South African populations. An atlas of 30 chromosome-level, fully-phased nuclear haplotypes from six USA isolates and nine Australian isolates was generated to capture the genomic composition of key oat crown rust lineages. This study provides evidence of nuclear exchange and recombination of haplotypes in both the USA and Australian *Pca* populations as mechanisms for the introduction of genetic diversity. The limitations of assuming clonal evolutionary history from virulence phenotyping is demonstrated by the detection of either sexual or cryptic genetic recombination in the *Pca* Australian population. Overall, the characterization of intercontinental migration of *Pca* at the haplotype level provides further impetus for molecular monitoring of rust pathogen populations on a global scale.

## Introduction

Rust fungi are a diverse group of biotrophic plant pathogens belonging to the Basidiomycete order Pucciniales (Bauer *et al*., 2006; Aime & McTaggart, 2020). With over 7,000 species and diverse hosts, these organisms cause many significant agricultural diseases (Kirk *et al*., 2008). Notably, many rust fungal species such as *Puccinia coronata* f. sp. *avenae* (*Pca* - oat crown rust), *P. graminis* f. sp. *tritici* (*Pgt –* wheat stem rust), *P. striiformis* f. sp. *tritici* (*Pst* – wheat stripe rust), and *P. triticina* (*Pt* – wheat leaf rust) propagate as dikaryons (two haploid nuclei) while infecting their cereal hosts, which has complicated the identification of virulence factors and genomics informed surveillance (Petersen, 1974; Figueroa *et al*., 2020).

The flax – flax rust (*Melampsora lini*) pathosystem was the basis for Flor’s pioneering work in plant pathology establishing the gene-for-gene model (Dodds, 2023), in which recognition of pathogen avirulence (Avr) effectors by plant resistance (*R*) proteins triggers signalling cascades and ultimately leads to cell death, an effective defence strategy against biotrophs (Jones & Dangl, 2006; Dodds & Rathjen, 2010; Chen *et al*., 2022; Dodds, 2023). However, understanding virulence evolution and host adaptation in rust populations has been hampered by their complex dikaryotic genome structures. The development of key molecular and bioinformatics technologies has resolved challenges posed by the ploidy changes in rust fungi through their complex lifecycles (Figueroa *et al*., 2022). Long-read sequencing and haplotype-aware assembly software enabled the first partial dissection of haplotypes to uncover high heterozygosity between nuclei of *Pca* and *Pst,* which highlighted the importance of nuclear phasing to create accurate rust genome references (Miller *et al*., 2018; Schwessinger *et al*., 2018). Further improvements in chromatin contact sequencing and phasing pipelines for rusts delivered the first chromosome-level and nuclear phased genomes for *Pgt*, *Pt*, and *Pca* (Li *et al*., 2019; Duan *et al*., 2022; Henningsen *et al*., 2022). In *Pgt*, this revealed that whole nuclear exchange without recombination precipitated the emergence of the devastating wheat stem rust Ug99 lineage (Li *et al*., 2019), leading to a paradigm shift that rust epidemiology must consider the movement of entire haplotypes. This has been made possible by the advent of high-fidelity long-read sequencing technology allowing the assembly of more and larger nuclear-phased rust genome references (Duan *et al*., 2022). This has been explored recently in *Pt*, where extensive somatic hybridization and intercontinental migration was uncovered through comparative genomics (Sperschneider *et al*., 2023).

Although molecular resources for studying rust fungi continue to grow, well-established areas of research for other plant pathogens remain poorly characterized in these pathogens. For instance, about 90% of Basidiomycota require different mating types to mate (heterothallic) (Kües *et al*., 2011), with compatibility controlled by two mating type loci, *a* and *b*. These loci encode a pheromone and receptor at the *a* (*PR*) locus and homeodomain transcription factor subunits at the *b* (*HD*) locus (Bölker *et al*., 1992; Burkhard *et al*., 1990). Heterothallic fungi are further divided into two main types; bipolar and tetrapolar, in which the *a* and *b* loci are linked and unlinked, respectively. The presence and location of conserved mating type genes in the rust species *Pgt*, *Pt*, and *Pst* suggests heterothallic and tetrapolar systems (Cuomo *et al*., 2017; Holden *et al*., 2023), although the role of these presumed mating type loci has not been functionally validated in rust fungi (Cuomo *et al*., 2017). It has been proposed that this system also plays a role in somatic combability for nuclear exchanges to occur between strains (Flor, 1964).

Oat crown rust disease caused by *Pca* results in significant yield losses worldwide and is difficult to control due to rapid host adaptation by the pathogen (Nazareno *et al*., 2018). Molecular evidence shows that sexuality is a clear driver of this rapid virulence evolution in USA populations and influences population structure in Sweden (Berlin *et al*., 2018; Miller *et al*., 2020; Hewitt *et al*., 2023). However, asexual *Pca* populations have also been observed to gain virulence following *R* gene deployment (Nazareno *et al*., 2018; Figueroa *et al*., 2023). Like for other rusts, key aspects of *Pca* biology and epidemiology remain unclear.

The role of somatic hybridization in *Pca* virulence evolution cannot be assessed without haplotype-phased genomes (Li *et al*., 2019; Sperschneider *et al*., 2023). There is one fully phased, chromosome level assembly (Pca203) and two partially haplotype-separated reference genomes from USA isolates (12SD80 and 12NC29) (Miller *et al*., 2018; Henningsen *et al*., 2022). Given the strong genetic differentiation between USA and South African isolates detected by Hewitt *et al*. (2023), the existing references are unlikely to capture the genetic variation of *Pca* globally. Thus, there exists ripe opportunity to address key questions for rust biology in *Pca* by further expanding global resources for the pathosystem that allow genomic comparisons at the haplotype-level. This study combines a haplotype-aware pangenome with the extensive collection of publicly available short-read data for *Pca* to further our understanding of the biology and epidemiology of *Pca* through the application of contemporary genomic-based comparative approaches.

## Materials and Methods

### Pca isolates, amplification, and phenotyping

Australian isolates were collected from 2020 to 2023 across six states and territories (ACT – Australian Capital Territory; NSW - New South Wales; QLD – Queensland; SA – South Australia; VIC – Victoria; WA – Western Australia). Isolates from Taiwan (TW) were collected from the College of Bioresources and Agriculture Experimental Farm at the National Taiwan University in 2020 and 2021 (Taipei, Taiwan). Oat lines for phenotyping were sourced from existing CSIRO seed stocks and the Australian Grains Genebank (AGG) (Henningsen *et al*., 2024).

Differences between infection procedures for reviving field samples, single pustule purification, and phenotyping are detailed in full in Methods S1. In all infections, susceptible oat seedlings were grown at 23C° for 16 hours light and 18C° for 8 hours dark. Plants were treated with 15 mL maleic hydrazide per pot at 9 days of growth immediately prior to inoculation. Inoculations were conducted using urediniospores in an oil or talc suspension and infected plants were kept in humidity chambers (90-99% relative humidity) for two days before removal to growth chambers under the same conditions as before. For phenotyping, infection types were recorded at 10-11 days post inoculation (dpi), with the final score chosen to reflect the most prevalent infection type across the biological replicates of the same differential line.

### DNA extraction and sequencing

DNA extractions for Illumina sequencing were performed with the G-Biosciences OmniPrep Genomic DNA isolation kit using 20-40 mg of rust spores as input. Libraries were generated using either the Illumina DNA PCR-Free Prep or the IDT Prism library preparation protocol depending on sample needs. Libraries were sequenced to 15-30X depth, 150 bp paired end reads with Illumina Novaseq by Azenta or the Australian Genome Research Facility (AGRF) (Table S1).

Extractions for high molecular weight (HMW) DNA from rust spores were completed as described previously by Li *et al*. (2019). HMW DNA was used for PacBio HiFi library preparation and sequencing to 30-50X depth at Azenta Life Sciences (formerly Genewiz) through Phase Genomics (US isolates) or AGRF (Australian isolates) (Table S2). Isolates 21ACT116, 20WA94, and 21WA134 were sequenced to 100-200X coverage due to high yield from a single lane of PacBio Revio.

Spores for Hi-C were prepared as described by Sperschneider *et al*. (2023) and detailed in Methods S2. Hi-C libraries were prepared at Phase Genomics, Seattle WA USA and sequenced with Illumina Novaseq by Azenta Life Sciences (formerly Genewiz) or were prepared and sequenced with Illumina Novaseq at the Ramaciotti Centre for Genomics, NSW Australia (Methods S2).

### Genome assembly, phasing, scaffolding, and annotation

Some contamination was apparent in the high-coverage HiFi read data for 20WA89, 21ACT116, and 21WA134 based on large and highly fragmented initial assemblies, so these reads were filtered with mash v2.0 (Ondov *et al*., 2019) as described in Methods S3. Raw reads for 20WA94 and 21WA139 and cleaned reads for 20WA89, 21ACT116, and 21WA134 were filtered on length with a 10 kb cutoff and were then randomly down-sampled with seqkit (v2.7.0) sample to 30-50X coverage (Shen *et al*., 2016). PacBio HiFi reads were assembled using Hifiasm with Hi-C integration and resulting contigs were cleaned, phased, scaffolded, and annotated according to Sperschneider *et al*. (2023) and described in detail in Methods S4. BUSCO values for each haplotype were determined using compleasm v0.2.1 (Huang & Li, 2023) with the odb10 Basidiomycota lineage dataset (Manni *et al*., 2021).

### Identification of putative mating type genes and population screening

Protein sequences for *STE3.2.1*, *STE3.2.2*, *STE3.2.3*, *bW-HD1* and *bE-HD2* from *Pgt* were BLASTed (tblastn) against the Pca203 genome reference with BLAST+ v2.13.0 (Camacho *et al*., 2009; Cuomo *et al*., 2017). Proteins overlapping the best hits were compared to the *Pgt* alleles and examined for conserved domains known to be present and intron/exon count before use in searching the other 30 haplotypes (Kües *et al*., 2011; Cuomo *et al*., 2017; Methods S4). Protein sequences for *STE3.2*, *bW-HD1*, and *bE-HD2* alleles were aligned with CLUSTALW, and phylogenetic trees were constructed with the built-in ‘raxml-bootstrap’ option (Larkin *et al*., 2007). Protein trees were rooted on alleles from *Pgt* CRL 75-36-700-3 (*Pgt STE3.2.1* and *Pgt bW-HD1*) with R package ‘ape’ and visualized with iTOL (Cuomo *et al*., 2017; Paradis & Schliep, 2019; Letunic & Bork, 2021).

To screen the *Pca* population for the presence of the *STE3.2* and *HD* alleles, the 3’ to 3’ DNA sequence of the *HD* locus and 5’ to 3’ sequences of *STE3.2.2* and *STE3.2.3* were extracted from each haplotype. These loci were sketched with mash v2.0 (-s 1000; Ondov *et al*., 2019) and then screened with the Illumina reads for 352 isolates (Miller *et al*., 2018, 2020; Henningsen *et al*., 2022; Hewitt *et al*., 2023; Ho *et al*., 2024; Table S1). Alleles were considered contained at or above 99% *k*-mer identity and 94% shared *k*-mers.

To compare the repetitive regions on chromosome 9, haplotypes were aligned with minimap2 v2.22 (-k19 -w19 -m200 -DP -r1000) and visualized with ggplot (Li, 2018; Wickham, 2016).

### Population screening for haplotype containment

The 352 *Pca* isolates with Illumina data (Table S1) were screened against the 32 available *Pca* haplotypes using mash v2.0 (Ondov *et al*., 2019). These haplotypes were processed with mash sketch (-s 100000) and mash screen was run comparing the Illumina data against the haplotype sketch file. Isolates with *k*-mer identity of ≥ 99.98 % and shared *k*-mers ≥ 99.65% were considered as likely containing the screened haplotype. Candidate hybrids were identified for having high identity to only one haplotype, or as having high identity to two haplotypes which were not contained within the same reference isolate.

### Phylogenetic analysis

Illumina reads for 352 isolates (Table S1) were mapped to the diploid assemblies for isolates Pca203, 90TX52, 20NSW19, 20QLD86, 20WA94, and 21ACT116 using bwa-mem2 v2.2.1 (Vasimuddin *et al*., 2019). Variants were called against the diploid assembly of Pca203 and individual haplotypes (3, 4, 5, 6, 13, 14, 23, 24, 25, 26), using freebayes v1.3.6 (--use-best-n-alleles 6) (Garrison & Marth, 2012). Variants were filtered with vcflib v1.0.1 (-f "QUAL > 20 & QUAL / AO > 10 & SAF > 0 & SAR > 0 & RPR > 1 & RPL > 1 & AC > 0") and vcftools v0.1.16 (--min-alleles 2 --max-alleles 2 -- max-missing 0.9 --maf 0.05) (Danecek *et al*., 2011; Garrison *et al*., 2022). Variants were converted to PHYLIP format using vcf2phylip (Ortiz, 2019). RAxML v8.2.12 was used to construct the Maximum Likelihood (ML) tree with 500 bootstraps (-f a -m GTRCAT -# 500 --no-bfgs) (Stamatakis, 2014). The resulting ML tree was visualized with iTOL (Letunic & Bork, 2021).

Variants called against the Pca203 complete reference for the 137 Australian isolates were filtered to include only biallelic single-nucleotide polymorphisms (SNPs). Variants were converted to PHYLIP format with vcf2phylip (Ortiz, 2019), and heterozygous calls were converted to missing (N) resulting in 391,118 SNPs. Splitstree CE v.6.2.1-beta was used to calculate Hamming distances and generate a neighbor net with 301 splits (Bryant & Huson, 2023; https://github.com/husonlab/splitstree6). The network was evaluated with the phi test for recombination in Splitstree CE v6.2.1-beta (Bruen *et al*., 2005).

### Comparison of pathotype clustering and phylogenetic tree structures

Infection types were converted to a binary system of 0 (avirulent, infection types 0 to 2) and 1 (virulent, infection types 3 to 4) and clustered with R v4.3.2 hclust() before conversion to a dendrogram (R Core Team, 2023). The Pca203 ML tree was pruned to contain only phenotyped Australian isolates with iTOL (Letunic & Bork, 2021) and was modified to be binary and ultrametric, with R package ‘ape’ before conversion to a dendrogram (Paradis & Schliep, 2019). The two dendrograms were compared with untangle_step_rotate_1side() from ‘dendextend’, with the phenotype clustering dendrogram being rotated to find the best structural match to the RAxML-derived dendrogram (Galili, 2015). The comparison was visualized with tanglegram(rank_branches = TRUE) from ‘dendextend’ to visualize topology without considering branch lengths (Galili, 2015). Finally, the two dendrograms were compared with cor_bakers_gamma() from ‘dendextend’ to calculate the correlation between the tree structures (Galili, 2015).

### Haplotype comparisons

Nucmer and DNAdiff from mummer v4.0.0 were run to determine the identity between all nuclear haplotype pairs (Kurtz *et al*., 2004; Marçais *et al*., 2018). Haplotypes were aligned with D-Genies v1.5.0 and visualized with ggplot2 (Wickham, 2016; Cabanettes & Klopp, 2018). The cactus-pangenome pipeline from cactus v2.6.6 was run on all haplotypes and variants were called for each haplotype as the reference (Hickey *et al*., 2023). The full procedure for determining shared haplotype blocks is detailed in Methods S5. Briefly, variant counts were binned and a cutoff of <= 50 non-ref SNPs per 100kb was applied. Adjacent shared regions for each sample-reference pair were merged, subjected to an ordered bedtools (v2.31.1; Quinlan & Hall, 2010) subtraction, and regions <= 50 Kb were removed before visualization.

## Results

### Contextualization of Australian and Taiwanese Pca isolates within the global population

From 2020 to 2023, 137 Australian *Pca* isolates were collected through community submissions across six Australian states and territories (ACT = 9, NSW = 36, QLD = 9, WA = 55, SA = 10, VIC = 18), and isolates were recovered from both wild (*n*=111) and cultivated oats (*n*=26) (Table S1). Furthermore, four Taiwanese isolates were collected from cultivated oat in 2020 and 2021. Whole genome sequence (WGS) data (Illumina) was generated from genomic DNA from all isolates (42X average genome coverage; Table S1). We generated a ML phylogenetic tree using the WGS data of all isolates and including published data of 211 isolates from the USA, South Africa, and Taiwan (total *n*=352) (Miller *et al*., 2018, 2020; Henningsen *et al*., 2022; Hewitt *et al*., 2023; Ho *et al*., 2024). Reads were mapped to both haplotypes of the Pca203 genome reference (Henningsen *et al*., 2022) and filtered variants (*n*=376,646) were used to construct the ML phylogenetic tree.

A total of 18 lineages were detected in the Australian *Pca* collection, which is unexpected as the population is proposed to consist of only four asexually-reproducing (clonal) genetic groups (Fig. **2a**; Park *et al*., 2022). Although clonality is influential in Australia as shown by the size and temporal persistence of the largest lineages (L18 – 48 individuals, L1 – 35 individuals), nine lineages consist of only one isolate. Further, only seven lineages were sampled across multiple years (4 years – L1, L9, L18; 3 years – L2, L11; 2 years – L5, L16), suggesting there are more lineages to be sampled (Fig. **2a**). Lineage diversity was greatest in WA (*n*=15), followed by SA (*n*=5), NSW/ACT (*n*=4), VIC (*n*=3), and QLD (*n*=2; Fig. **2b**). The five most abundant lineages (L1, L3, L9, L11, L18) were sampled on both wild and cultivated oat, suggesting that the populations on these different host types are not separate (Table S1). Only one lineage was detected from the four Taiwanese isolates, which are clones of isolate NTU-01.

**Figure 1.**
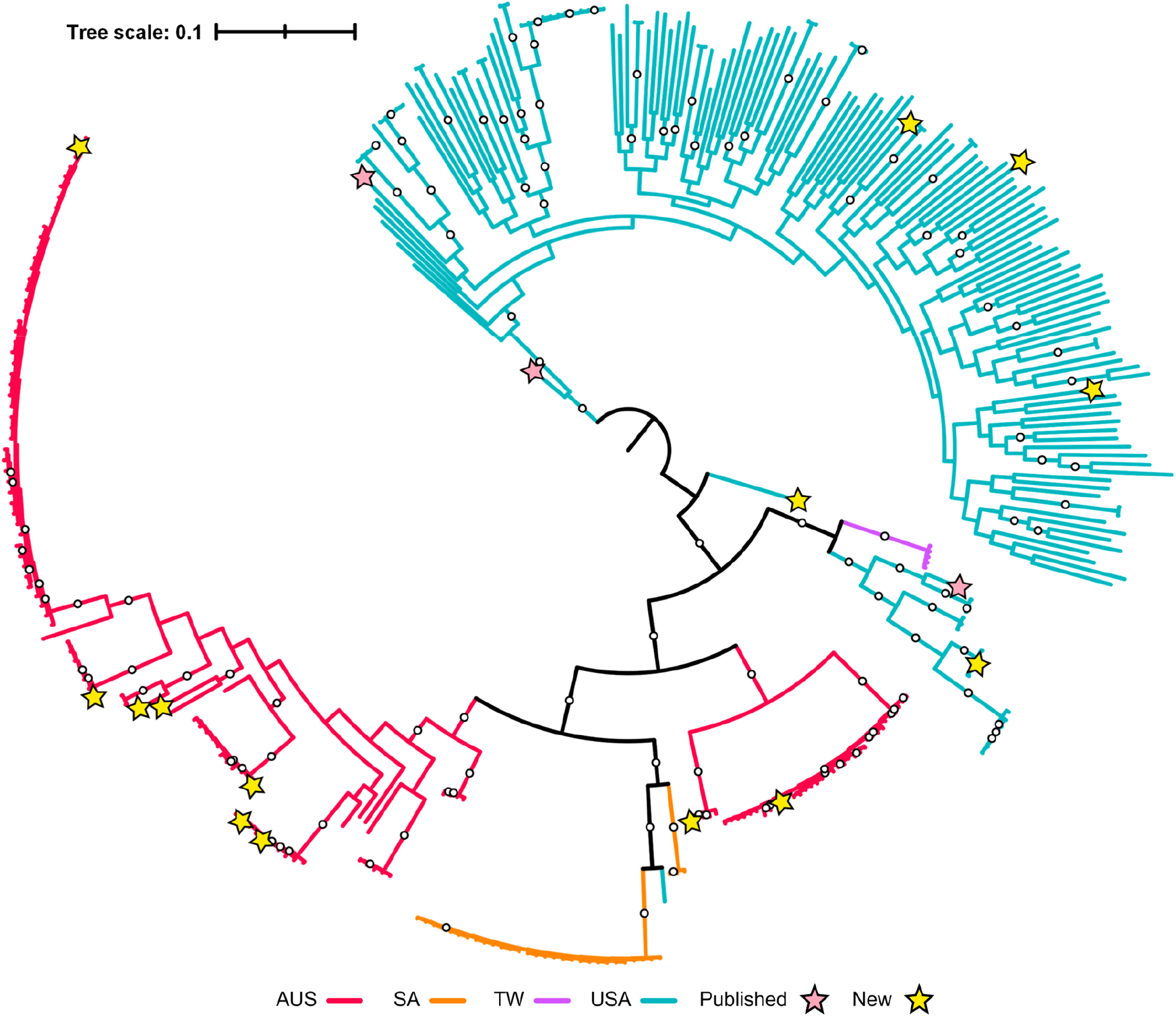
Midpoint rooted ML phylogenetic tree constructed by mapping reads from 352 *Puccinia coronata* f. sp. *avenae* (*Pca*) isolates and calling variants against the full Pca203 reference (hap1, hap2, and unplaced contigs). A total of 376,646 biallelic SNPs and 500 bootstraps were used for this analysis. Tree branches are colored by country of origin: AUS = Australia; SA = South Africa; TW = Taiwan; USA = United States of America. Bootstrap values above 80% shown as white circles. Yellow stars indicate isolates chosen for the haplotype atlas and red stars indicate isolates with existing references. Tree scale is mean substitutions per site.

**Figure 2.**
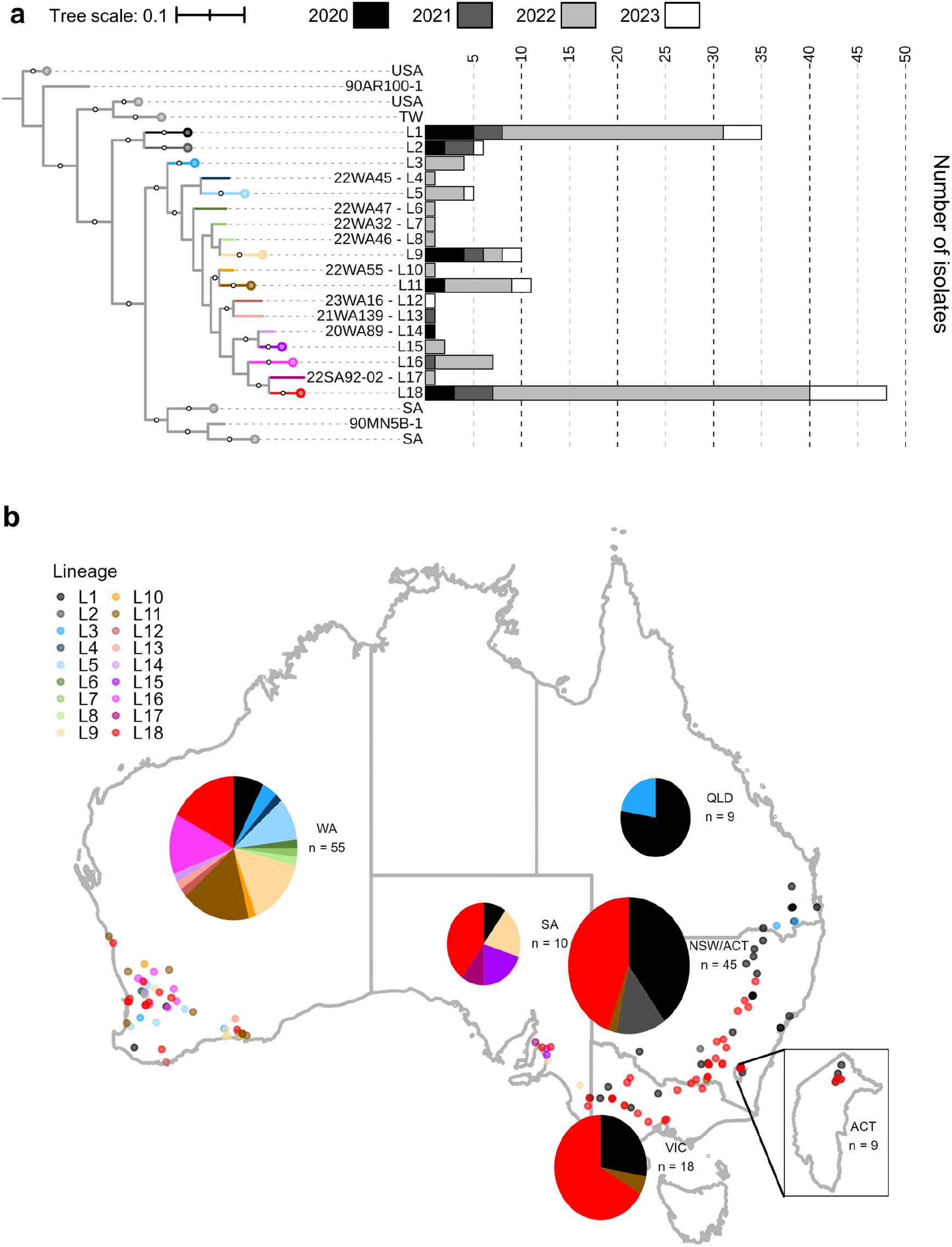
**a)** View of Australian lineages with clones collapsed in the midpoint rooted ML phylogenetic tree constructed for 352 *Puccinia coronata* f. sp. *avenae* (*Pca*) isolates based on 376,646 biallelic SNPs called against the full Pca203 reference. Bootstrap values (500 cycles) above 80% shown as white circles. Tree scale is mean substitutions per site. Stacked barplots show the number of isolates of each lineage collected in 2020 (black), 2021 (dark grey), 2022 (light grey), and 2023 (white) across Australian lineages. Tree branches for Australian lineage L1-L18 are colored by lineage to match points and pie charts in panel **b)** Distribution of *Pca* isolates and lineages across Australia. Pie charts are scaled on a logarithmic scale according to the total number of *Pca* isolates collected in their respective states. Pie chart for NSW/ACT includes isolates from both regions due to the small area of the ACT relative to NSW. Colors reflect lineage assignments.

To assess the robustness of virulence phenotypes as the basis for lineage assignment within the Australian *Pca* population, 95 of the 137 sequenced *Pca* isolates were tested on 27 genotypes from the Australian oat crown rust differential set (Henningsen *et al*., 2024; Fig. S2; Table S3). Most isolates in lineage L1 (20ACT25, 22NSW08) displayed broad virulence across the differential lines, while others had virulence to fewer lines (22QLD110, 22NSW107) (Fig. S2). Lineage L2 had the least virulence, with 21ACT116 and 23NSW13 only defeating resistance for two lines, Swan (carrying *Pc1*) and Pc70 (also known as H547). The remaining lineages (L3-L18) share many virulence traits. For example, isolates 22WA31 and 22WA54 from lineages L11 and L5 had nearly identical reactions. Clustering isolates by these phenotypic results is inconsistent with the phylogenetic relationships (Baker’s Gamma = 0.64, Fig. S3). Thus, virulence phenotypes (pathotypes) are poor predictors of lineage relationships in the Australian population.

### Construction of a haplotype-aware pangenome of Pca

From the phylogenetic analysis, we selected representative *Pca* isolates to generate haplotype-phased chromosome-level genome references that capture genetic diversity in Australia (*n*=9) and USA (*n*=6) (Fig. **S4a,b**). References were generated from eight Australian clonal groups (20QLD86 – L1, 21ACT116 – L2, 20WA72 and 20WA95 – L9, 20WA94 – L11, 21WA139 – L13, 20WA89 – L14, 21WA134 – L16, 20NSW19 – L18), with lineage L9 sampled twice (20WA72, 20WA95) as a baseline for comparing near-identical haplotypes (Fig. **S4a**). We also chose three contemporary buckthorn-derived isolates (18MNBT34, 18MNBT36, 18MNBT50) from the USA sexual population (Hewitt *et al*., 2023; Fig. **S4b**) and three isolates from USA clonal lineages (90TX52, 90AR100, 90MN4B; Miller *et al*., 2018; Fig. **S4a,b**).

We assembled nuclear phased genome references of 15 isolates following established computational workflows using hifiasm with incorporated Hi-C chromatin contact data (Duan *et al*., 2022; Henningsen *et al*., 2022; Sperschneider *et al*., 2023), resulting in 30 complete nuclear haplotypes (hap3-hap32; Fig. **1**, see yellow stars; Fig. **3a**; Table S2). Nuclearphaser detected only five phase swaps (in three isolates) that required correction. Four within-haplotype mis-assemblies were also corrected in hap5, hap9, hap10, and hap19. Scaffolding resulted in 18 chromosomes in each haplotype (chr1 to chr18; Fig. **3b**). An average of 84.44% of *trans* and 96.75% of *cis* and *trans* Hi-C contacts occurred on chromosomes within the same haplotype which reflects accurate haplotype separation and nuclear assignment. *Pca* haplotypes were on average 98.4 Mb long with 42.7% repeat coverage and 19,700 gene annotations (Table S4). Haplotype BUSCO completeness averaged 94.85% with less than 2% duplication, which is comparable to haplotype-phased assemblies for *Pgt* and *Pt* (Table S4; Li *et al*., 2019; Duan *et al*., 2022; Sperschneider *et al*., 2023). Haplotypes were consistent for most genome characteristics (Table S4). Chromosome sizes were likewise consistent except for chr9, which harbours a large repetitive region of approximately 0.1 to 1.4 Mb (Fig. **3b,c**). Most assemblies break within this locus, so the exact size of the region is not certain.

**Figure 3.**
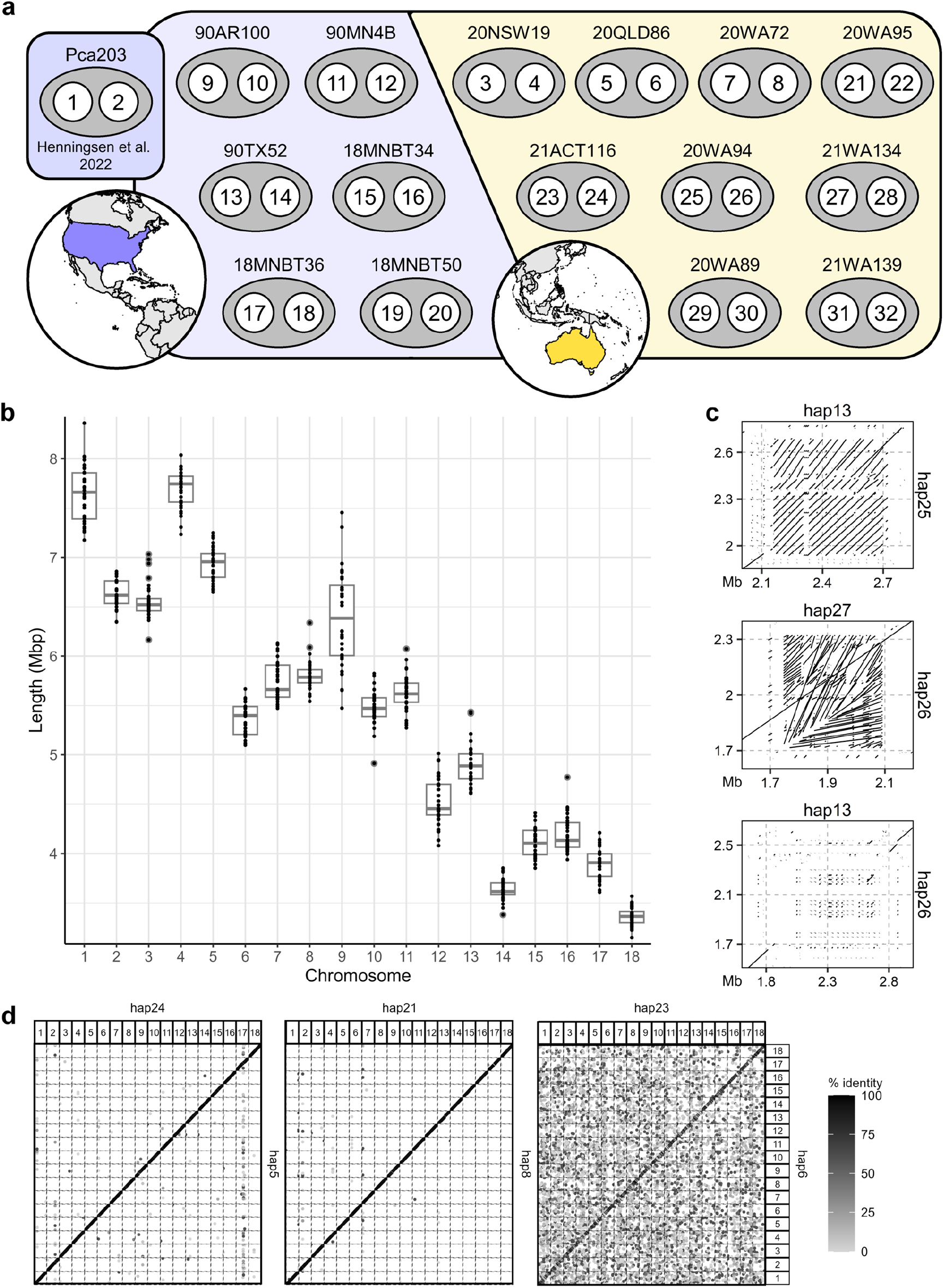
**a**) Haplotype numbers assigned to nuclear genotypes of 15 *Puccinia coronata* f. sp. *avenae* (*Pca*) isolates from the USA (purple) and Australia (yellow), in addition to previously published haplotypes from Pca203 (1 = “A”; 2 = “B”). **b**) Boxplots of chromosome lengths for the 32 *Pca* haplotypes with black dots indicating chromosome size of each haplotype. Box top and bottom boundaries are the upper and lower quartiles, respectively. Lines within boxes represent the mean. Lines extending below and above boxes delimit minimum and maximum values. **c**) In descending order: alignments of chromosome 9 regions containing STE3.2 genes between hap13 and hap25 (*STE3.2.2*), hap26 and hap27 (*STE3.2.3*), and hap13 and hap26 containing different *STE3.2* alleles. **d**) Dotplot of genome alignment between *Puccinia coronata* f. sp. *avenae* (*Pca*) haplotypes hap5 and hap24 (left), hap8 and hap21 (middle) and hap6 and hap23 (right). Chromosomes are numbered along the x- and y-axis. Greyscale indicates percent identity along alignment.

Pairwise haplotype comparisons validated the clonal relationship between 20WA72 and 20WA95 from L9, as their matching haplotypes (hap7 ≈ hap22; hap8 ≈ hap21) had very high sequence identity (99.98%). In contrast, comparison of the two haplotype sequences within any isolate showed high levels of divergence and heterozygosity (98.73 - 99.45% identity), with most inter-isolate haplotype comparisons similarly diverse (98.65 - 99.98% identity). The only exception to this was the haplotype pair hap5 (20QLD86 from L1) and hap24 (21ACT116 from L2) which showed high 99.94% sequence identity (99.78-99.86% alignment coverage, 9,228 SNPs) comparable to haplotypes in the clonal 20WA72 and 20WA95 isolates (99.44-99.97% alignment coverage, 99.98% identity, 1,051-1,701 SNPs; Fig. **3d**). As the other haplotypes from 20QLD86 (hap6) and 21ACT116 (hap23) were extremely dissimilar (98.39% identity, 96.17-97.85% alignment coverage, 481,662 SNPs), it is likely that lineages L1 and L2 are related through somatic hybridization and nuclear exchange (Fig. **3d**).

### Characterization of mating loci in the Pca pangenome and overall population

To better understand the role of putative *a* and *b* mating type loci in *Pca*, we characterized the *bW-HD1/bE-HD2* and *STE3.2* loci using sequences identified in *Pgt* (Cuomo *et al*., 2017). Pca203 has one copy of *STE3.2.3* on chr9A and one copy of *STE3.2.2* on chr9B. Similarly, all isolates in the *Pca* pangenome carry one near-identical copy each of *STE3.2.2* and *STE3.2.3* on opposite haplotypes of chromosome 9 (Fig. S**5a,b**). The *mfa* pheromone precursors were also identified close to these *STE3.2* alleles (Fig. **S5a**). The related gene *STE3.2.1* was invariant and present in a single copy on chr1 in all haplotypes except in hap11 (90MN4B) which had two copies, supporting previous findings that it is unlikely to be involved in mating compatibility (Cuomo *et al*., 2017). The *STE3.2.2* and *STE3.2.3* alleles are adjacent to the large repetitive region on chr9 which is allele-specific, explaining the unusual chr9 length distribution (Fig. **3b**, Fig. **S5c,d**). We then characterized the *bW-HD1 bE-HD2* (*HD*) allele pairs across haplotypes in the pangenome. All isolates contained different *HD* alleles in each haplotype on chr4 and a total of 13 alleles were recorded (Fig. **S5e,f**). Using these characterized loci, we screened Illumina short reads from the entire *Pca* collection (*n*=352) and found that all isolates contain both *STE3.2.2* and *STE3.2.3* alleles and two different alleles at the *HD* locus (Fig. **S6a**, Table S5, Table S6). In *Pca* isolates where one or both *HD* alleles were not characterized in the pangenome, short read mapping still indicates heterozygosity at the *HD* locus (Fig. S7). The observed genotype frequency for both the *PR* and *HD* loci in the USA sexual population (*n*=88 isolates derived as aecia from the sexual host buckthorn in the UMN nursery) are 0% homozygous and 100% heterozygous (*PR* locus alleles *n*=2, *HD* locus alleles *n*=11). This is a significant departure from the expected 50:50 (homozygous:heterozygous) ratio for the biallelic *PR* locus in Hardy-Weinberg equilibrium (*p* = 6.55 x 10^-21^). Likewise, the *HD* locus genotype ratio is significantly different than expected (12% homozygous, 88% heterozygous; *p* = 5.70 x 10^-4^). This suggests that both loci contribute to mating type compatibility resulting in exclusive heterozygosity, meaning that *Pca* likely has a tetrapolar mating system given that the *PR* and *HD* loci are on separate chromosomes.

### Pca isolates from the USA, Australia, and Taiwan are related through somatic hybridization and migration

To assess the role of somatic hybridization in the evolution of *Pca* we employed *k*-mer containment, phylogenetics, and comparative genomics approaches used to identify somatic hybridization events in other rust species (Li *et al*., 2019; Sperschneider *et al*., 2023). In our analysis, *k*-mer identity of ≥ 99.98 % and shared *k*-mers ≥ 99.65% accurately identified clones of the *Pca* reference isolates. For example, L18 isolate 20NSW20 has high *k*-mer containment for hap3 and hap4 from the L18 reference 20NSW19 (100% *k*-mer identity and ≥ 99.93% shared *k*-mers). As established in *Pgt* and *Pt*, phylogenetic trees constructed from an isolate’s entire genome show clonal lineages as discrete clades, while single-haplotype trees merge lineages that share the reference haplotype into a single clade (Li *et al*., 2019; Sperschneider *et al*., 2023). Multiple putative nuclear exchange events were identified across the entire *Pca* dataset using these criteria (Table S1; Table S7).

The full haplotype comparisons we described earlier are the strongest evidence for somatic hybridization between L1 and L2 and *k*-mer containment and phylogenetic tree results also support this hypothesis. L1 isolates had high containment for hap5, hap6, and hap24 (> 99.98% identity, > 99.75% shared *k*-mers), but not hap23 (< 99.86% identity, < 96.99% shared *k*-mers) (Fig. **4a**; Table S7). Conversely, L2 isolates had high containment for hap23, hap24, and hap5 (> 99.98% identity, > 99.82% shared *k*-mers) and low containment for hap6 (< 99.64% identity, < 92.50% shared *k*-mers) (Fig. **4a**; Table S7). Further, trees constructed from these haplotypes individually support hybridization between L1 and L2. Phylogenies generated from hap5 and hap24 both show L1 and L2 isolates as members of a single clade, while those for hap6 and hap23 show L1 and L2 as discrete clades (Fig. **4b-e**). These analyses also suggest a somatic hybridization event linking Australia and Taiwan, implicating direct or indirect migration of *Pca* (Fig. **4f**). The five Taiwanese isolates (L-TW) have high *k*-mer containment for hap6 (99.99% identity, > 99.84% shared *k*-mers) but not hap5 (< 99.70% identity, < 93.65% shared *k*-mers) (Fig. **4a**; Table S7). This is also supported by the phylogenetic trees, as L1 and L-TW isolates comprise separate clades in the hap5 tree but cluster together in the hap6 tree (Fig. **4b,d**). Since we did not sample the other haplotype for L-TW in the current pangenome, we cannot infer the history of haplotype exchange.

**Figure 4.**
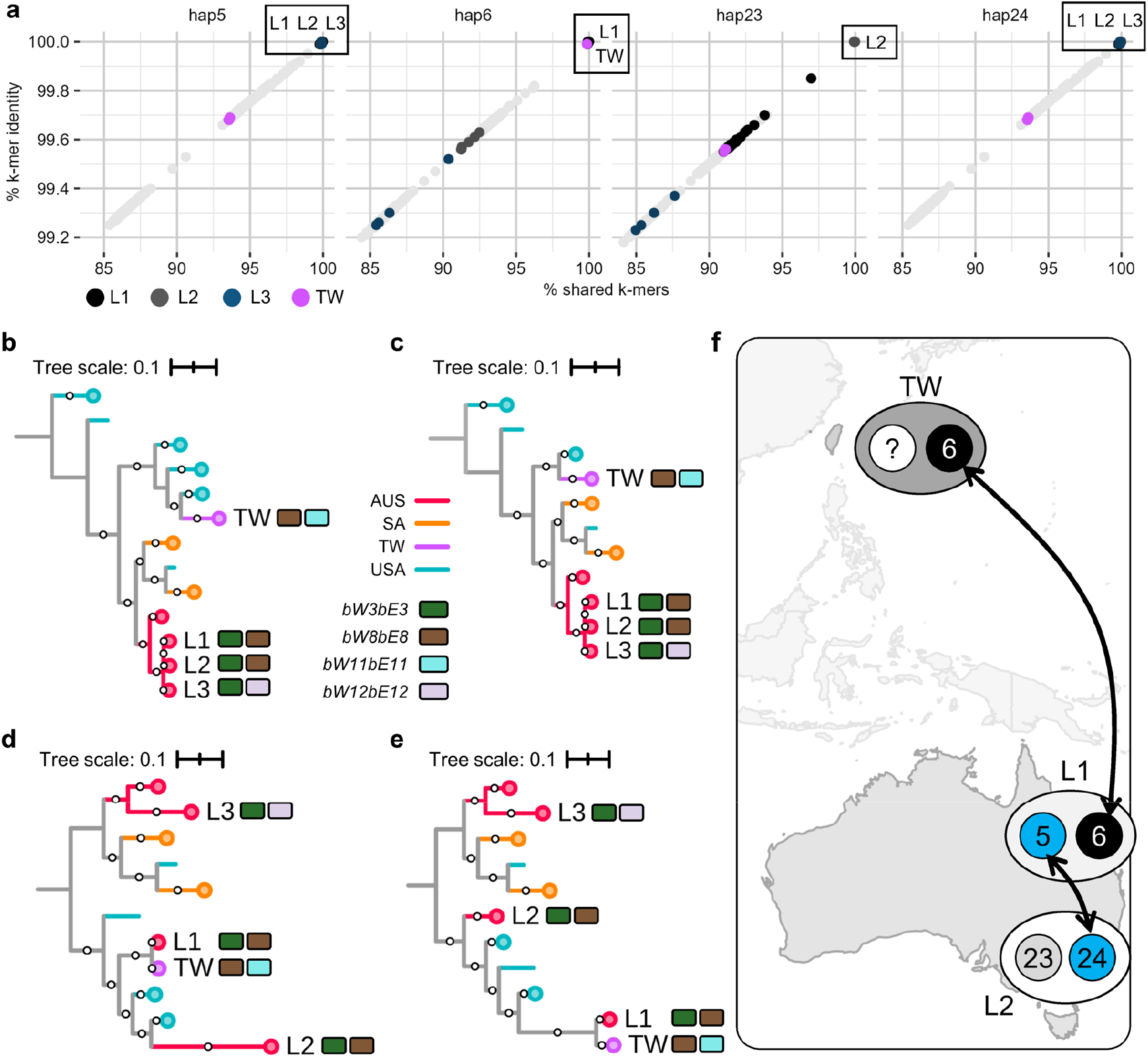
**a)** Plot of % shared *k*-mers (x-axis) versus % *k*-mer identity (y-axis) for 352 *Pca* isolates on hap5, hap6, hap23, and hap24. Colors indicate relevant lineages (L1-L3, TW =Taiwan); light grey points are all other isolates. **b-e)** Maximum likelihood trees for **b)** hap5 (225,727 SNPs) **c)** hap24 (239,896 SNPs) **d)** hap6 (198,467 SNPs) and **e)** hap23 (191,316 SNPs). Tree branches are colored by country of origin: AUS = Australia; SA = South Africa; TW = Taiwan; USA = United States of America. 500 bootstraps were generated for each tree and bootstraps 80% and higher are shown as white circles on branches. Collapsed branches are indicated with circles at leaf tips. Mating type alleles for hybrid lineages are shown as rectangles to the right of branches. Tree scales are mean substitutions per site. **f)** Proposed relationships between *Pca* lineages and haplotypes overlaid on a map of Oceania and Asia.

The *k*-mer containment and phylogenetic analyses also support the occurrence of somatic hybridization and nuclear exchange within the USA *Pca* population. Members of the 90TX52 lineage (L-1990) have high containment for both 90TX52 haplotypes (100% identity, > 99.92% shared *k*-mers) (Fig. **S8a**; Table S7). Seven clonal isolates collected in 2017 (L-2017) from the southern USA (LA, TX, FL) contain hap14 (100% identity, > 99.89% shared *k*-mers), but not hap13 (< 99.85% identity, < 96.65% shared *k*-mers) (Fig. **S8a**; Table S7). Phylogenetic trees constructed separately from hap13 and hap14 SNPs further support somatic hybridization between L-1990 and L-2017, as they appear as one clade in the hap14 phylogeny and separate clades in the hap13 phylogeny (Fig. **S8b,c**). As the second haplotype of the 2017 isolates was not sampled in the current pangenome, we cannot establish the history of hap14 inheritance (e.g., donor in the hybridization event) (Fig. **S8d**).

Mating allele composition of putative hybrid isolates agrees with the postulated nuclear exchange events, as somatic hybrid isolates have the same *HD* locus allele as the haplotype they are proposed to contain (Fig. **4b-e**; Fig. **S8b,c**; Table S5). The assemblies for hap5 and hap24 likewise have the same alleles at both mating loci (*STE3.2.2* and *bW3bE3*; Fig. S5). The other pangenome haplotypes do not appear to be related through somatic hybridization events to *Pca* isolates in the existing collection. However, 90MN14B and 90MN7B were identified as likely clones of 90MN4B (100% identity, > 99.92% shared *k*-mers) for which short-read data is not available (Table S7). Altogether, our results support the occurrence of somatic hybridization in the broad *Pca* population.

### The Australian and USA Pca populations are shaped by genetic recombination

The *k*-mer containment analysis also yielded some results that were incompatible with the hypothesis of nuclear exchanges. For instance, L18 isolates had high containment for hap3 and hap4 (L18) and hap25 (L11) (100% identity, > 99.91% shared *k*-mers) but not hap26 (L11) (< 99.94% identity, < 98.56% shared *k*-mers; Table S7). However, L11 isolates only had high *k*-mer containment for hap25 and hap26 (100% identity, > 99.95% shared *k*-mers) but neither hap3 nor hap4 (< 99.95% identity, < 98.70% shared *k*-mers), as would be the case if L11 and L18 were somatic hybrids (Table S7). Haplotype-specific phylogenetic trees reflected the same relationships, with L18 and L11 isolates forming a single clade in the hap25 phylogeny and discrete clades in the hap3, hap4, and hap26 phylogenetic trees (Fig. **S9a-d**). Whole-haplotype alignments between hap3, hap4, and hap25 clarified this non-reciprocal relationship. Hap25 has high-identity alignments from hap3 and hap4 in large blocks covering >95% of this nuclear genome, with 0 to 3 breakpoints between hap3 and hap4 sequences per chromosome. This is consistent with the frequency of recombination breakpoints in a single meiotic event reported in other rust fungi (Fig. **S9e**; Anderson *et al*., 2016). As further confirmation, we obtained variants from a pangenome graph of all haplotypes and recovered the same recombination breakpoints as before from hap3 and hap4 SNP-sparse regions on hap25 (Fig. **S9f**). This is consistent with hap25 being derived as a haploid product of meiosis from lineage L18 as part of a sexual cross.

We next explored the contradictory finding that L3 isolates had high containment for haplotype hap5 (= hap24) as well as hap26 from 20WA94 (> 99.98% identity, > 99.84% shared *k*-mers), despite these all having the *bW3bE3 HD* allele pair (Fig. **4b**; Fig. S5; Table S5; Table S7). In support of the haplotype containment results, we found that L3 isolates cluster with L1 and L2 isolates in the hap5 and hap24 phylogenetic trees but with L11 in the hap26 tree, while these lineages all form discrete clades in the hap6, hap23, and hap25 phylogenetic trees (Fig. **4c-f**; Fig. **S9a,b**). The hap5/24 and hap26 haplotypes all contain the *bW3bE3 HD* allele, while L3 is heterozygous for b3/b12 alleles suggesting some recombination involved in the relationships between these isolates. Alignment of hap5 and hap26 again revealed numerous high-identity segments covering about 45% of haplotype genome, in a pattern that suggests recombination (Fig. **S10a**; Table S8). A model to explain these results is that L3 shares a nucleus (hap5) with lineage L1 and L2 through a nuclear exchange event, while hap26 in L11 is a meiotic recombinant between the two nuclei (hap5 and an unknown haplotype) in L3 (Fig. **S10b**). In this scenario, L11 results from a sexual cross between L3 and L18. The unique sequence amounting to approximately half of the unknown L3 haplotype should also contain *bW12bE12*.

Given the evidence for recombination in the hap25 and hap26 genomes, we evaluated the entire Australian *Pca* population for recombination by generating a splitstree network which showed reticulation between most Australian lineages. The *phi* test was significant (p < 10^-10^), suggesting that recombination has occurred to shape the extant Australian population. We next characterized haplotype recombination across the entire *Pca* pangenome using the pangenome graph-derived variants approach validated above in our parent-F1 comparison. A subset of six USA haplotypes were assessed first, which showed a pattern of small recombination blocks that reflects frequent sexual recombination in the USA facilitated by the prevalence of *R. cathartica* (Fig. **5a**). Further, each haplotype added only 4 to 9% coverage, with hap18 having approximately 30% coverage by the preceding five USA haplotypes. This suggests the full diversity of the USA sexual population has not been sampled in the haplotype pangenome.

**Figure 5.**
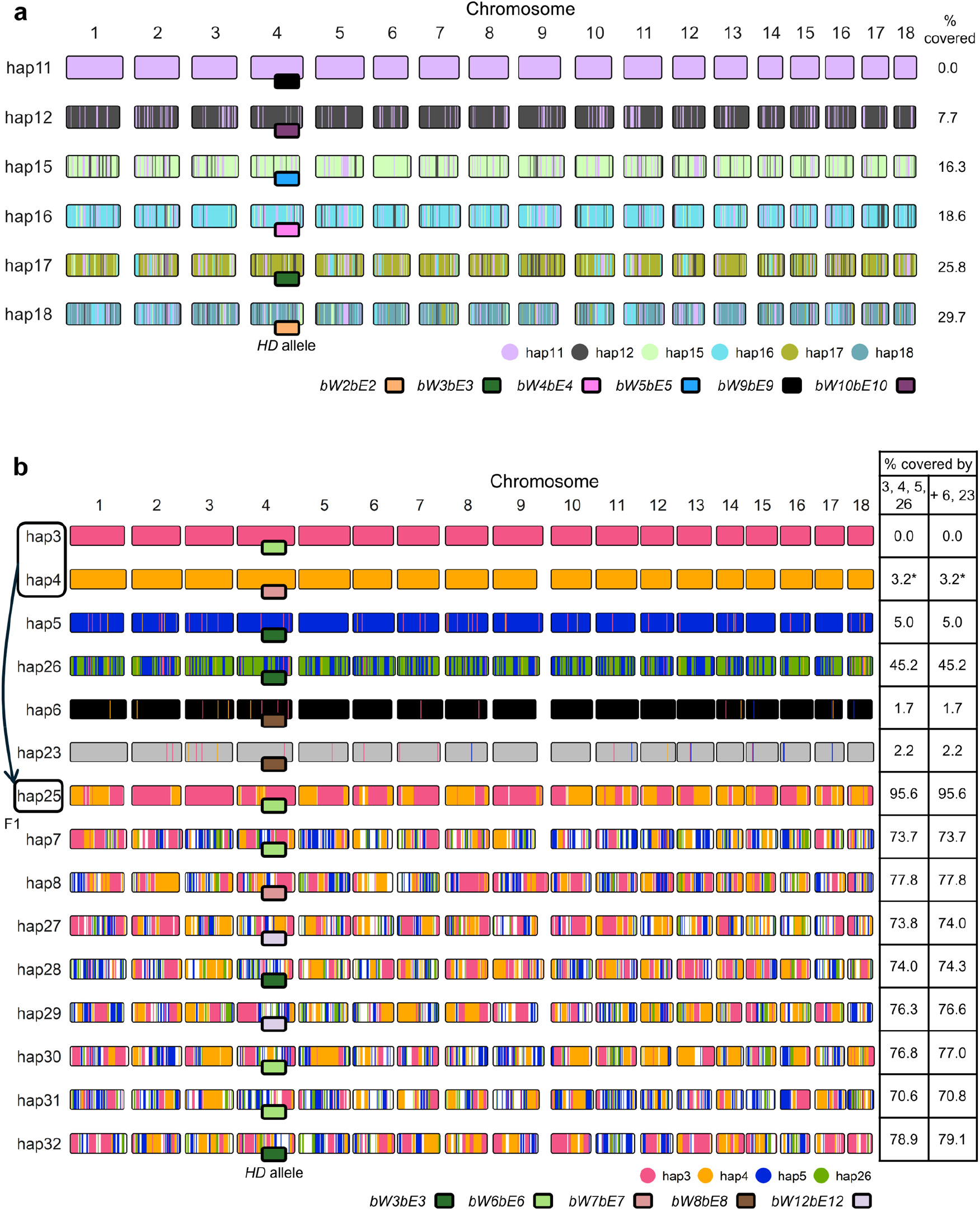
**a)** Shared recombination blocks across six *Puccinia coronata* f. sp. *avenae* (*Pca*) haplotypes identified by identifying 100 kb sequence bins with less than 50 non-reference variants hierarchically across hap11, hap12, hap15, hap16, hap17, and hap18. Shared sequences with hap11 were assigned across the other five haplotypes first, after which the unassigned areas of hap12 were evaluated on the four haplotypes below in the hierarchy, and so on. **b)** Shared haplotype blocks between 15 Australian *Pca* haplotypes from founder representatives hap3, hap4, hap5, hap26 and divergent haplotypes hap6 and hap23 generated in a similar manner to panel **a)** with hap3 being first in the hierarchy. The haplotype with the shorter name was kept as the representative in cases where haplotypes are nearly identical, as they show redundant information in this analysis (hap8, hap7, and hap5 shown; hap21, hap22, and hap24 not shown). Chromosome fill color represents unassigned regions or regions shared with haplotypes earlier in the hierarchy. Unassigned regions from hap7, hap8, hap27, hap28, hap29, hap30, hap31, and hap32 are filled white. Table on the right side shows percent coverage of each haplotype by those listed in each column. *HD* locus alleles are indicated with fill color of rectangles at the chromosome 4 midpoint. Asterisk (*) indicates that the haplotype blocks on hap4 from hap3 are not shown for visual clarity.

Among the Australian haplotypes, hap6 and hap23 are clearly divergent from the others, while all other haplotypes appear related through recombination (Fig. **5b**). Hap3 and hap4 have contributed to all recombinant haplotypes, while hap5 and hap26 have contributed to all except hap25. The large size of recombination blocks between Australian haplotypes relative to those shared between USA haplotypes suggests that recombination is relatively infrequent. However, Australian haplotypes in L3-L18 are approximately 75% covered by hap3, hap4, hap5, and hap26, consistent with founding by four haplotypes. Considering our earlier results with hap5 and hap26, we hypothesize that the remaining 25% is likely present in L3, which has not been represented in the pangenome yet. Therefore, Australian lineages L4-L17 may have been founded by limited recombination between the L3 and L18 lineages (Fig. **6a**). The divergent hap6 and hap23 haplotypes in L1 and L2 have not contributed to the L3-L18 recombining population.

**Figure 6.**
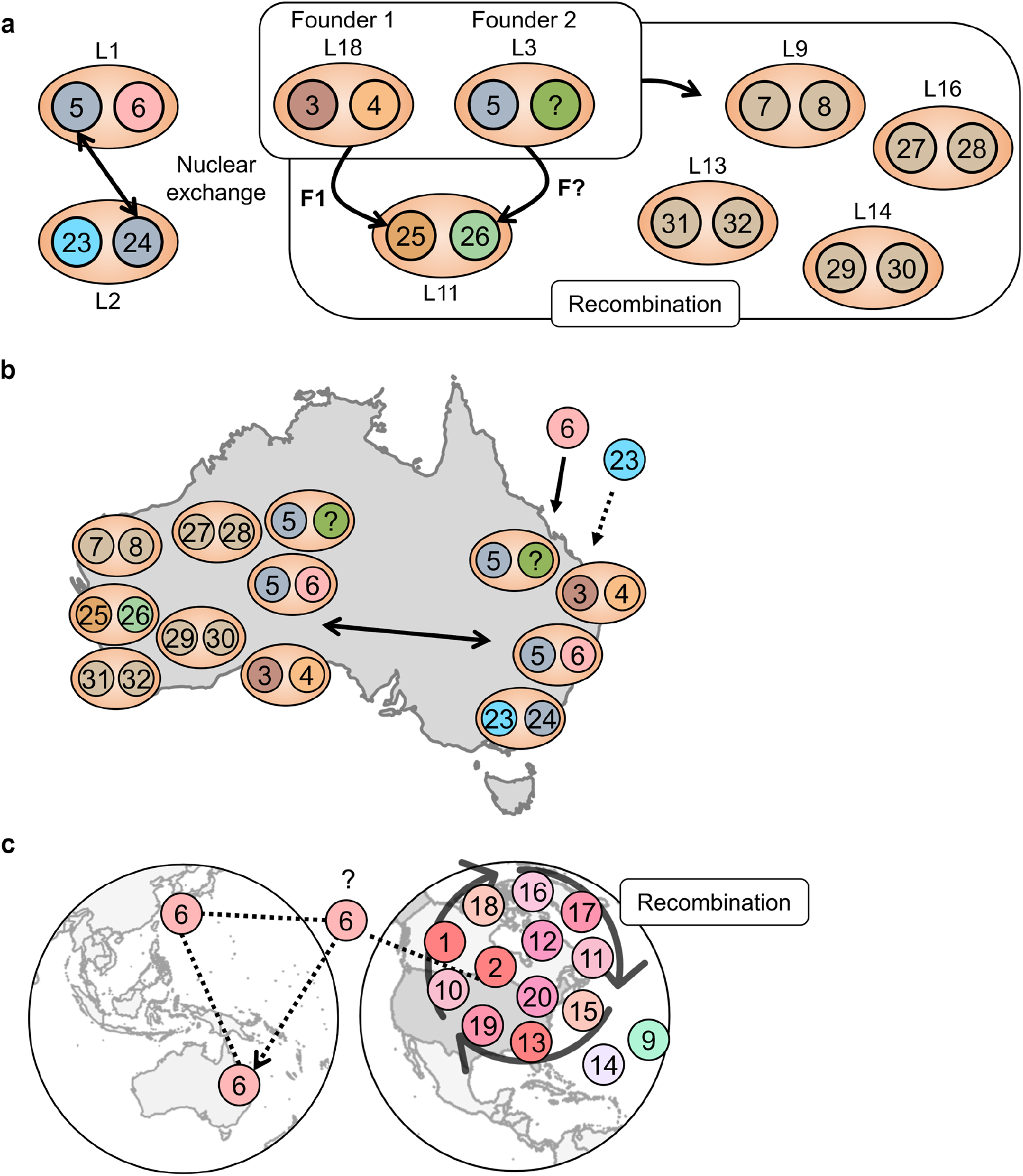
**a)** Diagram showing the proposed relationship of haplotypes in the Australian *Puccinia coronata* f. sp. *avenae* (*Pca*) population to the two postulated founder lineages L18 and L3. Four colors are assigned to founding haplotypes (brown – hap3, orange – hap4, blue – hap5, green – hap‘unknown’) and derivative haplotypes are color combinations of their ancestors. **b)** Known distribution of haplotypes in the Australian *Pca* collection in either Western Australia or eastern Australia (Australian Capital Territory, New South Wales, Queensland, South Australia, Victoria). **c)** Recombination between closely related USA *Pca* haplotypes (shades of pink) and hap6 with simplified possible pathways for hap6 migration.

Recombination analysis performed on all haplotypes demonstrated that the USA and Australian haplotypes overall have not had recent recombination, as the total amount of shared sequence between USA and Australian haplotypes was generally less than 5% (Table S8). The only exceptions were hap6 and hap23, which share approximately 50% and 15% of their sequence with the USA haplotypes, respectively (Fig. **S12b**). Hap1 and hap2 from Pca203 contribute most to hap6 (31.4%), while the other USA haplotypes share small regions (<5%) with the remaining hap6 sequence (Fig. **S12b**). Hap9 and hap14 are divergent from other USA haplotypes, but hap14 and hap6 have approximately 11% of their sequence in common (Table S8). The USA population is shaped by frequent recombination between diverse haplotypes, which is illustrated by hap20 having only 45% of its sequence represented by 11 other USA haplotypes (Fig. **S12b**). Historic isolate Pca203 clearly had an influential founding effect on the USA population, with hap1 and hap2 contributing from 21.3 to 57.6% of the sequence of other USA haplotypes (Fig. **S12b**; Table S8).

## Discussion

In this study we investigated the genetic relationships between newly sequenced Australian and Taiwanese *Pca* isolates and previously evaluated populations from the USA and South Africa (Miller *et al*., 2018, 2020; Hewitt *et al*., 2023). The rich genotypic diversity of this pathogen necessitated the construction of a haplotype-aware pangenome for the species to capture genotypic variability across different geographic areas. We focused on haplotypes from fifteen isolates representing Australian and USA *Pca* lineages as an initial step. This haplotype atlas allowed us to examine mechanisms that contribute to virulence evolution such as migration, somatic hybridization, and sexual recombination in *Pca* (Li *et al*., 2019; Figueroa *et al*., 2020; Sperschneider *et al*., 2023).

The role of somatic hybridization (nuclear exchange) in the evolution of dikaryotic rust fungi was proposed in the 1950-1960’s (Flor, 1964; Dodds, 2023) and was investigated in *Pca* under controlled inoculations and conditions (Bartos *et al*., 1969). Unfortunately, most early studies assessing somatic hybridization in cereal rust fungi failed to ensure that isolate cross-contamination did not explain detection of new races/virulence profiles. Indisputable proof of nuclear exchange in rust fungi was found by comparing the first chromosome level assembly of wheat stem rust *Pgt*21-0 and sequences from epidemic isolate *Pgt* Ug99 (Li *et al*., 2019), which showed that one nuclear haplotype of *Pgt*21-0 is nearly identical to one of the haplotypes of *Pgt* Ug99. The power of haplotype resolution to track the migration and nuclear exchanges shaping global rust populations was subsequently demonstrated for *Pt* (Sperschneider *et al*., 2023). This study provides evidence that somatic hybridization is also occurring in *Pca* populations in Australia, USA and possibly Taiwan, supporting that nuclear exchanges are common to rust fungi.

Newly acquired and existing whole-genome sequence short-read data and haplotype atlas also enabled population-level *in silico* characterization of mating type control in *Pca*. In other Basidiomycota such as smuts and mushrooms, mating type has been shown to maintain dikaryotic state, regulate life cycle, and enforce self/non-self recognition during mating (Nieuwenhuis *et al*., 2013; Coelho *et al*., 2017). Although the role of mating loci has not been functionally characterized in the rusts, our finding that all 353 *Pca* individuals in this study are heterozygous at both the *PR* and *HD* loci suggests that mating type controls critical biological functions in *Pca*. Consistent with the findings of Luo et al. (2024), pheromone precursor *mfa2* is linked closely to *STE3.2.2* in *Pca*; however, we identified a 55 amino acid *mfa3* pheromone precursor linked to *STE3.2.3* which was not reported in their study. Analysis of the *HD* alleles in the haplotype pangenome released by our study support most sequences imputed for *Pca HD* locus alleles published by Luo et al. (2024). There is one discrepancy between the *bW5* allele between the studies in which the first 12 N-terminal amino acids of *bW5* defined by Luo et al. (2024) are not encoded in the haplotypes used to define *bW5* in this study. In addition, the ‘*bW8bE8*’ allele pair reported by Luo et al. (2024) was only found in a single isolate (90MN5B) that was not included in our pangenome.

The *HD* locus configuration has been used in *Pt* and *Pst* to support conclusions regarding migration and population structure derived from genomic and phylogenetic approaches (Holden *et al*., 2023; Sperschneider *et al*., 2023). However, several divergent haplotypes (∼500-600K SNPs) included in the *Pca* pangenome have identical alleles at both mating type loci (i.e., hap6 and hap23, hap3 and hap13, hap5 and hap17). Thus, *HD* locus allele containment solely should not be used to infer lineage membership or haplotype composition.

We investigated the contribution of genetic recombination to the evolution of *Pca* in the USA and Australia. Small and frequent recombination blocks among USA haplotypes were detected and are consistent with widespread sexual reproduction enable by the prevalence of *R. cathartica* in North America (Miller *et al*., 2020; Hewitt *et al*., 2023). Prior to this study, molecular characterization of the Australian population was limited to 12 *Pca* isolates using DNA amplification fingerprinting and virulence phenotypes (Brake *et al*., 2001). Based on this, it was proposed the Australian *Pca* population was evolving by mutations in clonal lineages, despite the high phenotypic diversity of the pathogen.

We uncovered 18 lineages in the Australian *Pca* population, in contrast to the *Pca* collections from Taiwan and South Africa, which contain only one or two clonal lineages. This is a contrasting result to cereal rusts in Australia, like *Pgt* and *Pt*, which appear to be exclusively clonal, with the presence of only two and five lineages, respectively (Li *et al*., 2019; Sperschneider *et al*., 2023). The discovery of many lineages in the Australian *Pca* population was unexpected given its presumed asexuality (Park *et al*., 2022) and is evidence against mutations and incursions as the sole drivers of *Pca* evolution in the country. The best-described sexual host (*R. cathartica*) is not present in Australia or Oceania; however, other *Rhamnus* species (e.g., *R. alaternus, R. lycioides*, *R. palaestina*) have been reported as aecial hosts for *Pca* (Dinoor, 1962; Hemmami *et al*., 2006; Nazareno *et al*., 2018). Notably, *R. alaternus* was introduced to Australia as an ornamental and is currently managed as an invasive weed (Stajsic & VicFlora, 2023).

Both network analysis and whole-genome comparisons to define recombination blocks provided evidence of recombination among most Australian lineages, although infrequent when compared to the USA *Pca* population. It is yet unclear where or when these events occurred, and their rarity could be explained by sporadic access to a sexual host or uncommon parasexual events. Parasexuality akin to somatic hybridization that involves karyogamy and haploidization to generate recombinants without the aecial host is also proposed to occur in rust fungi (Harder, 1984), although this phenomenon has not been experimentally validated using molecular and genomic tools. The use of field collections hampers validation of parasexuality as isolates resulting from either somatic recombination or a sexual cross would be genotypically indistinguishable. The recombination among Australian lineages likely explains the high phenotypic diversity of *Pca* in Australia (Park *et al*., 2022), which was poorly correlated to lineage structure as has been reported in other *Pca* populations (Hewitt *et al*., 2023).

The splitstree network showed clear reticulation between lineages L3 through L18. Interestingly L3 and L18, which are positioned at opposite ends of the network, are also implicated in recombination events connected to L11. Hap25 is likely the meiotic product of hap3 and hap4, while hap26 may be a recombinant derived from L3 haplotypes (hap5 and unknown). Therefore, it seems that L3 and L18 could have undertaken a sexual cross to produce L11. While intercontinental migrations could explain these results, the likelihood of F1 haplotypes migrating together or in separate events is much lower than the hypothesis that rare sexual/parasexual cycles may occur. Together with the evidence of somatic hybridization in L1 and L2, we propose that the extant Australian *Pca* population was derived from three ancestral isolates. Two of these have undergone sexual and/or parasexual processes and the third donated nuclei via somatic nuclear exchange.

Our results suggest that it would be prudent to continue molecular monitoring of *Pca* and investigate the relationship between *R. alaternus* or other *Rhamnus* species and *Pca* in the Australian context, as the presence of these invasive species may present significant risk for virulence evolution to the detriment of the oat industry. Ongoing efforts to expand this haplotype-aware pangenome to include members from other continents will further our understanding of the evolution and global movement of *Pca*.

## Acknowledgements

We would like to thank Jakob Riddle (USDA-ARS) for his technical support. We would like to acknowledge the contribution of the Plant Pathogen ‘Omics Initiative consortium in the generation of data used in this publication. The Initiative is supported by funding from Bioplatforms Australia, enabled by the Commonwealth Government National Collaborative Research Infrastructure Strategy (NCRIS).

We also thank rust sample donors, with special thanks to Allan Rattey.

## Competing Interests

The authors declare no competing interests.

## Author Contributions

MF, PD, SFK, JS, BJS, and ES planned and supported the project. DL, ECH, Y-FH, and ESN amplified and prepared pathogen materials for sequencing and phenotyping. ECH and JS analysed the data. ECH, JS, MF, and PD prepared the manuscript and figures. All authors reviewed and edited the manuscript.

## Data Availability

Reference genomes are available in the CSIRO Data Access Portal (https://data.csiro.au/collection/csiro:61932). PacBio, HiC and genomic short reads are available on NCBI under BioProject PRJNA1063754. Scripts used to perform analyses and generate figures are available on GitHub (https://github.com/henni164/Pca_pangenome).

## Funding

This project was supported by the Bioplatforms Australia Plant Pathogen ‘Omics Initiative, GRDC project CSP2204 007RTX, USDA-NIFA BBSRC award 2022-67013-36505, the CSIRO Research Office, and grant 109-2313-B-002-028-MY3 from the National Science and Technology Council of Taiwan. ECH was supported by the ANU University Research Scholarship and ANU/CSIRO Digital Agriculture PhD Supplementary Scholarship.

## Supplementary Figures

**Figure S1.**
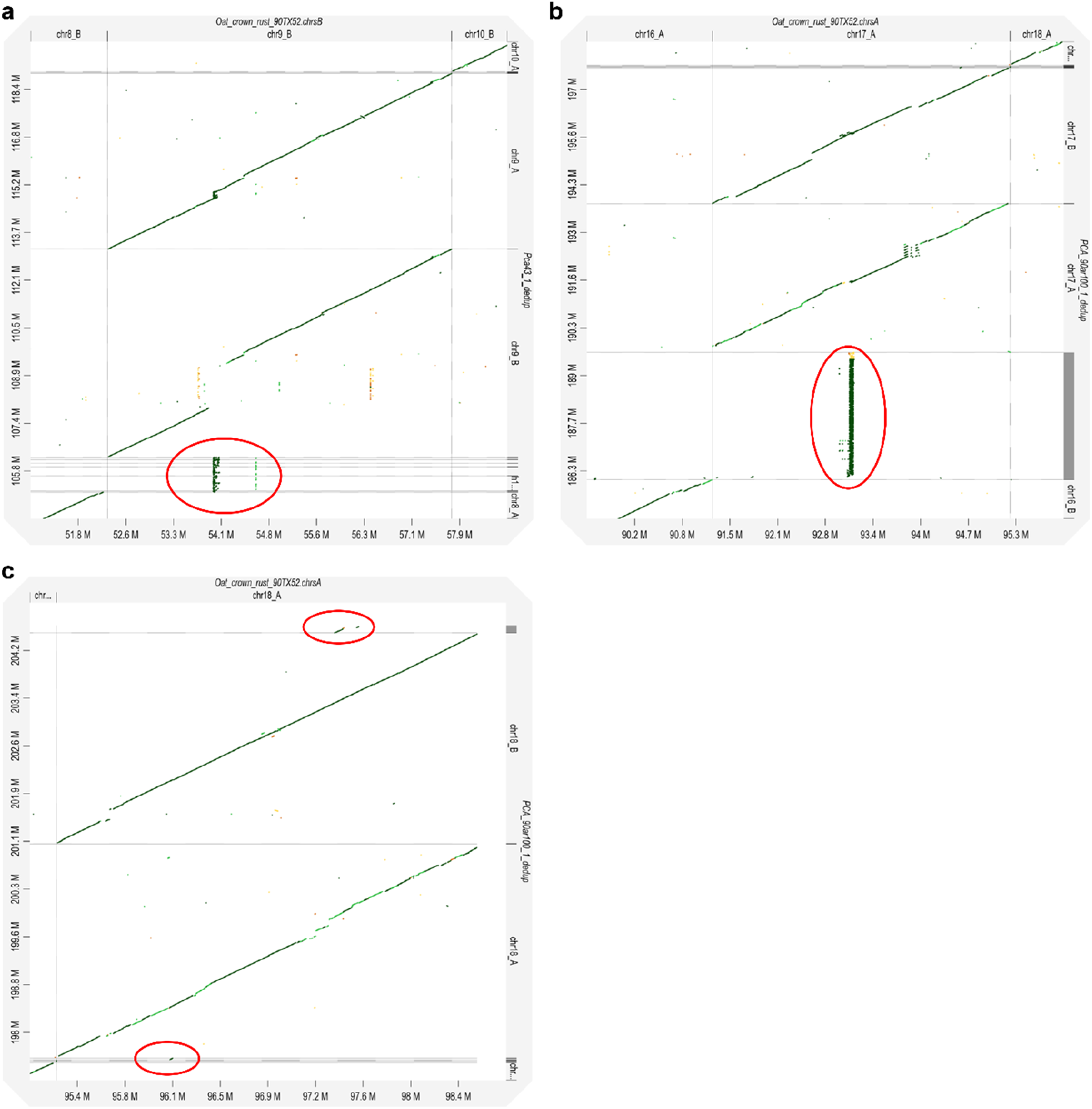
Examples of three main unplaced contig types: **a**) extra copies of chromosome 9 repeat-rich region **b**) extra copies of chromosome 17 ribosomal repeats **c**) extra copies of other genome sequences which are already represented in chromosomes.

**Figure S2.**
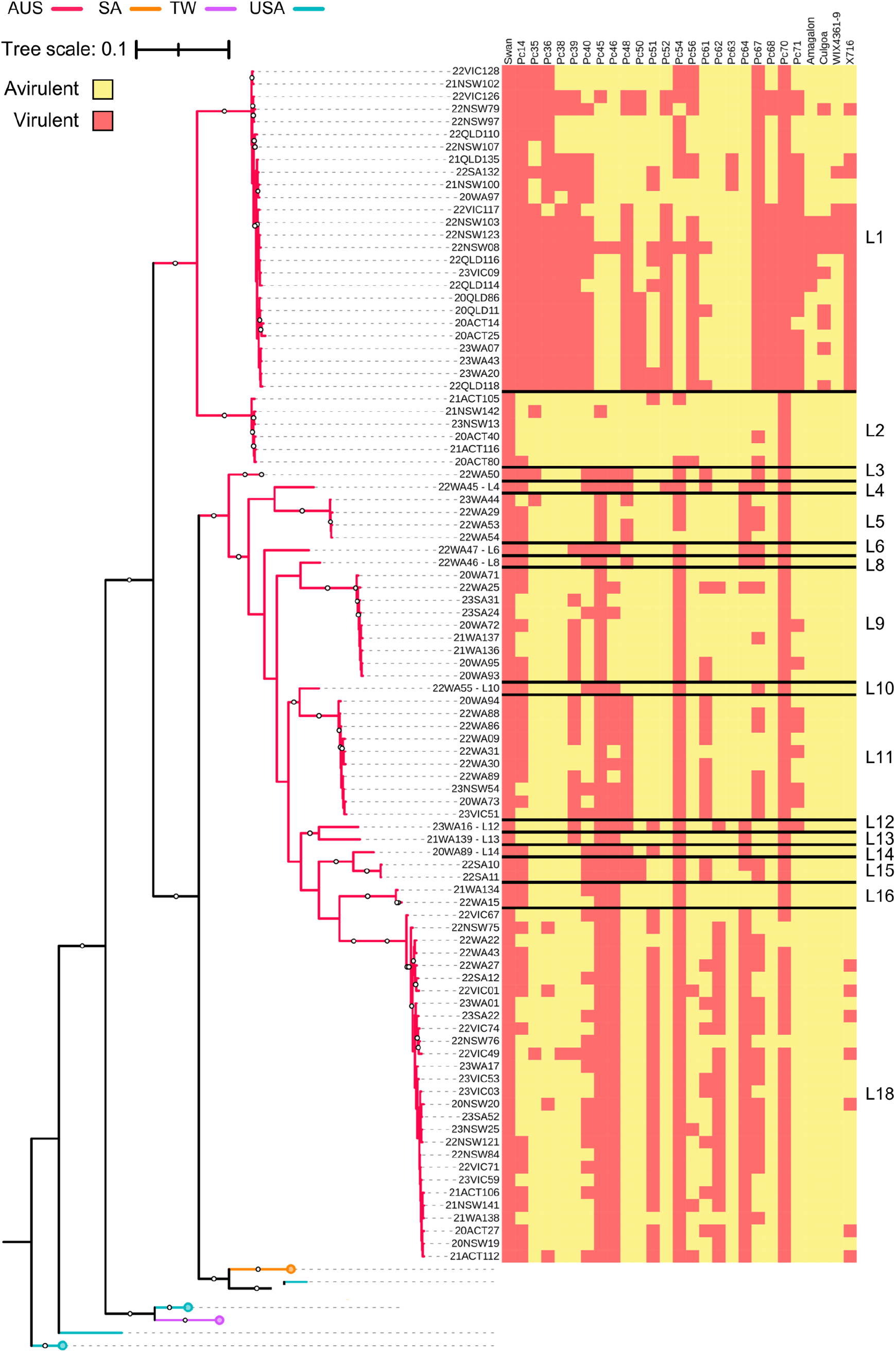
Phylogenetic tree constructed by mapping reads from 352 *Puccinia coronata* f. sp. *avenae* isolates to Pca203 hap1, and hap2, and unplaced contigs. 376,646 biallelic SNPs and 500 bootstraps were used to produce the maximum likelihood phylogenetic tree. Tree branches are colored by country of origin: AUS = Australia; SA = South Africa; TW = Taiwan; USA = United States of America. Bootstrap values are percentages (100 = 100%). Tree scale is mean substitutions per site. Heatmap represents isolate virulence on the differential lines (virulent = red, avirulent = yellow). Isolates that were not phenotyped were pruned from the tree, resulting in the removal of L7 and L17.

**Figure S3.**
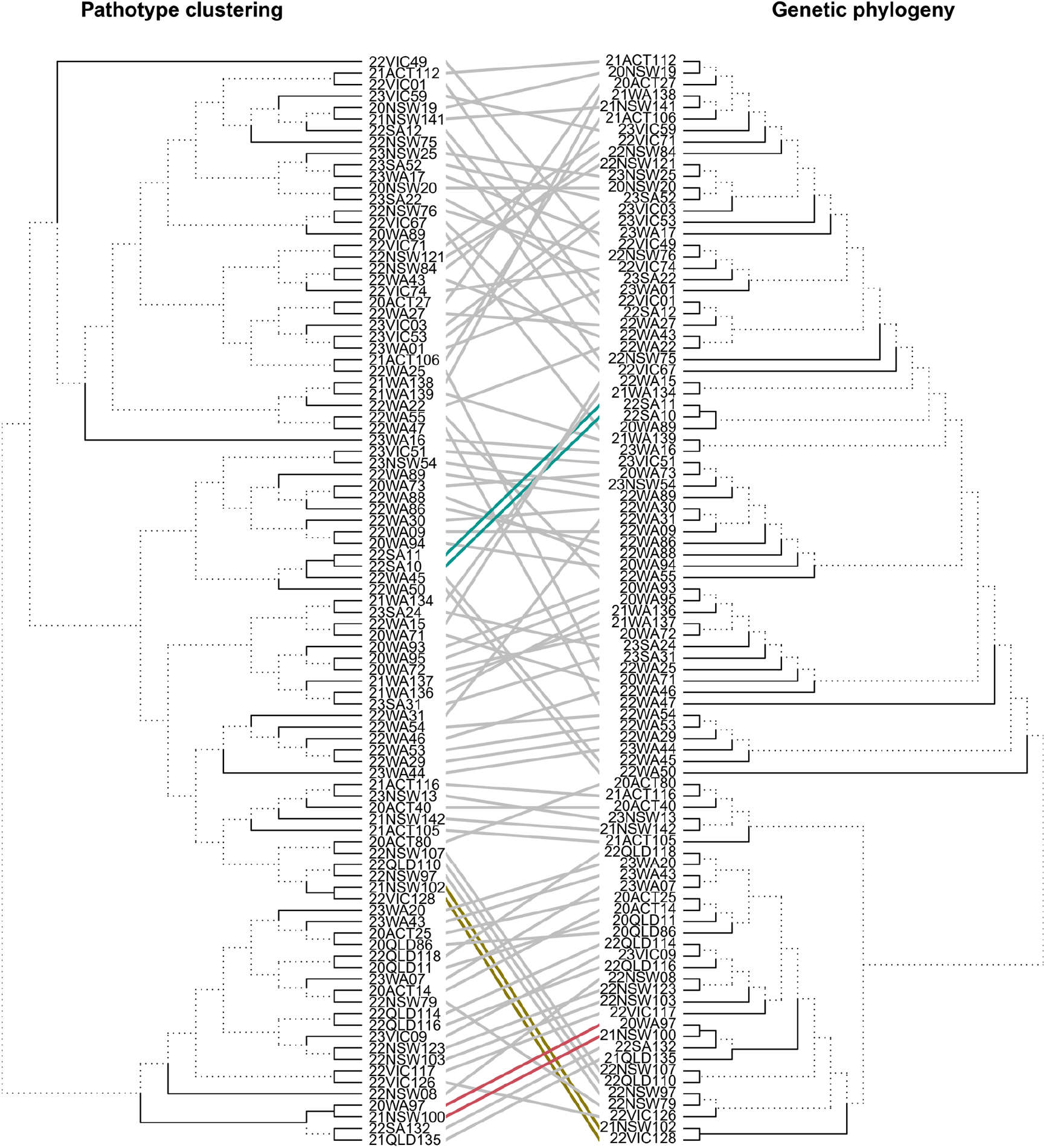
Comparison between tree topologies of 16 Australian *Puccinia coronata* f. sp. *avenae* lineages from clustering by pathotype (left) versus phylogenetic relationships (right). The ML tree (right) was generated with 376,646 SNPs from 352 *Pca* isolates against the complete Pca203 genome (hap1, and hap2, and unplaced contigs), which was pruned to contain only phenotyped Australian isolates and midpoint rooted. The R package ‘tanglegram’ was used to rotate the pathotype clustering tree branches until the best match to the phylogenetic tree was found. Branch lengths are arbitrary in the visualization and assessment of similarity. Solid black lines in the tree structure indicate edges found in both trees. Lines in the center connect the same isolate across trees, with colored lines showing clusters containing more than one isolate which are structurally identical.

**Figure S4.**
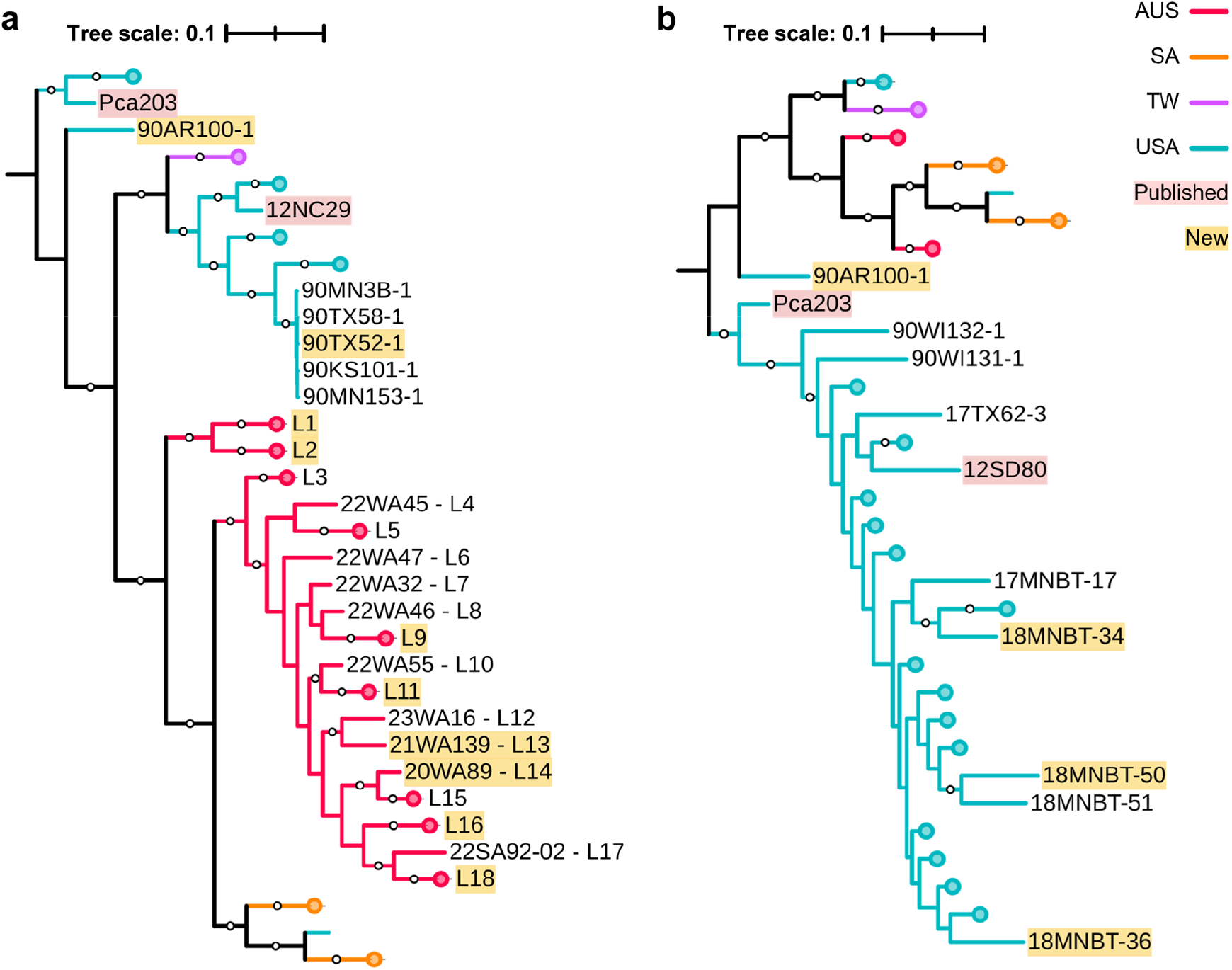
Midpoint rooted maximum likelihood phylogenetic tree of 352 *Puccinia coronata* f. sp. *avenae* (*Pca*) isolates constructed by mapping reads and calling variants against the full Pca203 reference (hap1, hap2, and unplaced contigs). 376,646 biallelic SNPs and 500 bootstraps were used. **a)** Collapsed tree view of primarily Australian lineages (L1 to L18). Except for Pca203, the first two digits of the name of the *Pca* isolate reflect year of collection, followed by state and sample identifier. **b)** Collapsed tree view showing USA lineages. USA *Pca* lineages were not numbered as the population is highly diverse. Tree branches are colored by country of origin: AUS = Australia; SA = South Africa; TW = Taiwan; USA = United States of America. Bootstrap values above 80% shown as white circles. Yellow labels indicate isolates chosen for the haplotype atlas and red labels indicate isolates with published references. Tree scales are mean substitutions per site.

**Figure S5.**
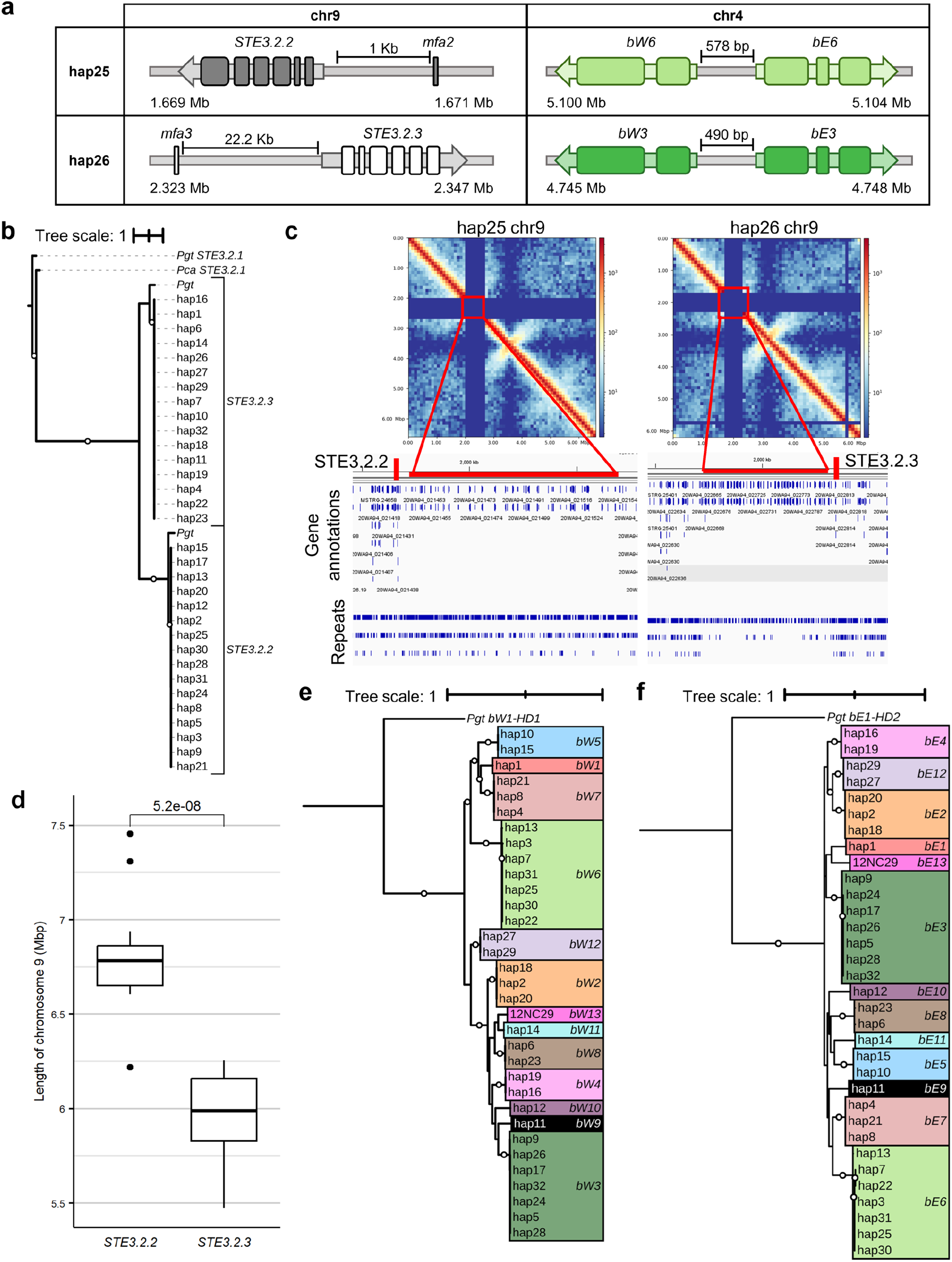
**a**) Orientation and arrangement of *STE3.2* and *mfa* (*PR*) alleles on chromosome 9 (chr9) and *HD* alleles on chromosome 4 (chr4) of hap25 and hap26 from *Puccinia coronata* f. sp. *avenae* (*Pca*) isolate 20WA94. **b)** Phylogenetic tree of *STE3.2* alleles in 32 *Pca* haplotypes, rooted at *Puccinia graminis* f. sp*. tritici* (*Pgt*) *STE3.2.1.* **c**) Diagram showing chromatin contact maps, and gene and annotations around the *STE3.2* locus (approximate position shown by labelled red vertical lines) on chromosome 9 in hap25 and hap26. Repetitive sequences adjacent to the *STE3.2* alleles correspond to multimapping reads resulting in loss of signal in the contact maps. **d**) Boxplots showing chromosome 9 length distribution separated by which *STE3.2* allele is present. **e**) Phylogenetic tree of *bW-HD1* alleles rooted at *Pgt bW1-HD1* and **f)** *bE-HD2* alleles rooted at *Pgt bE1-HD2* in 32 *Pca* haplotypes. Colors indicate identical *HD* alleles within each tree and allele pairs across trees (i.e. *bW3* and *bE3* are always found in the same haplotype), as recombination between *bW* and *bE* was not detected in our haplotype atlas. Branches with bootstraps over 80% are shown with white circles at the midpoint. Tree scales are mean substitutions per site.

**Figure S6.**
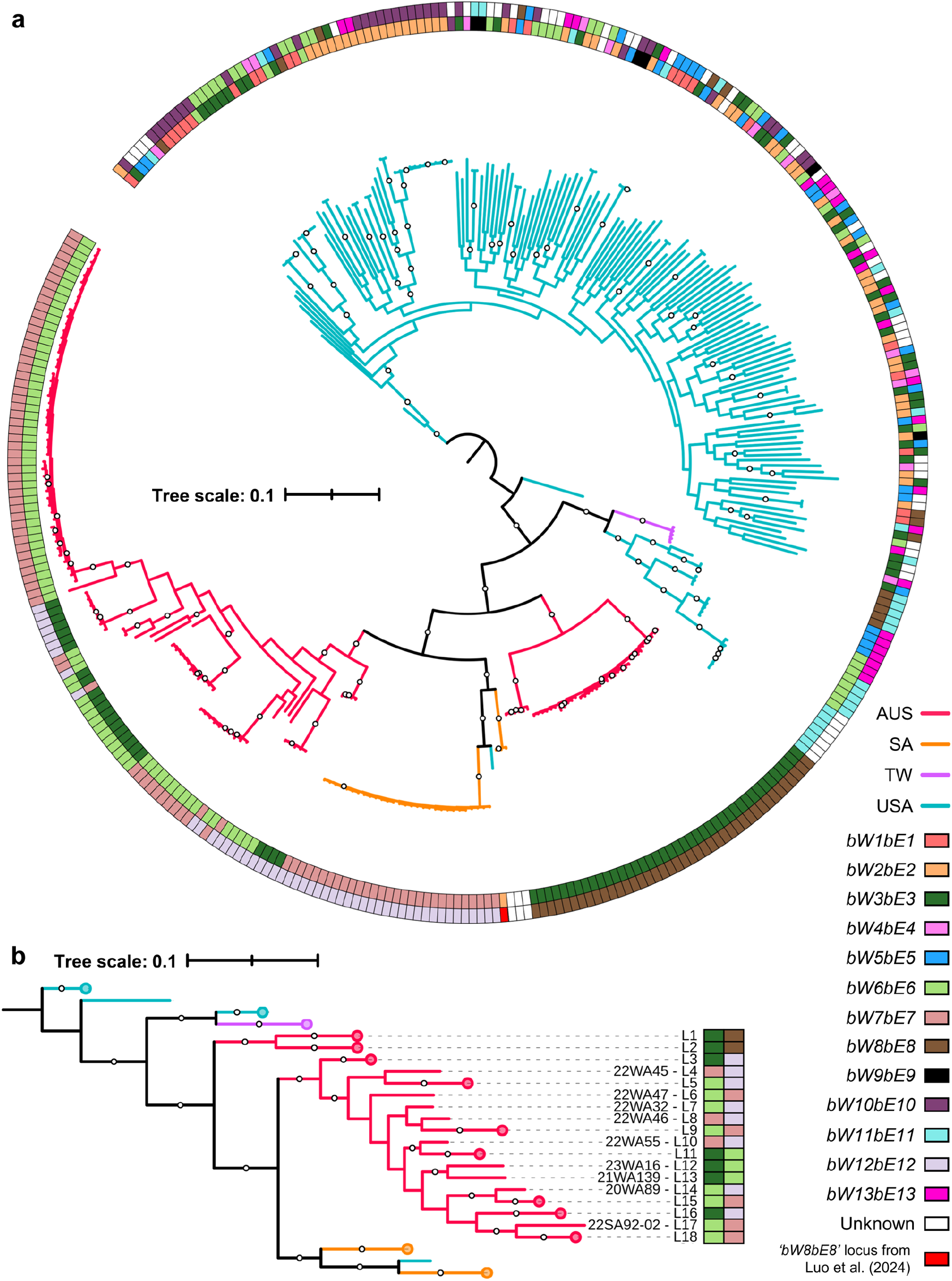
**A)** Midpoint rooted phylogenetic tree constructed by mapping short reads from 352 *Puccinia coronata* f. sp. *avenae* isolates and calling variants against the complete Pca203 genome (hap1, hap2, unplaced contigs). 376,646 biallelic SNPs were used over 500 bootstraps to produce the maximum likelihood tree. Bootstrap values are percentages (100 = 100%). **b)** Collapsed view showing Australian lineages L1-L18. *HD* alleles are shown as rectangles next to branches. Tree branches are colored by country of origin: AUS = Australia; SA = South Africa; TW = Taiwan; USA = United States of America. Tree scales are mean substitutions per site.

**Figure S7.**
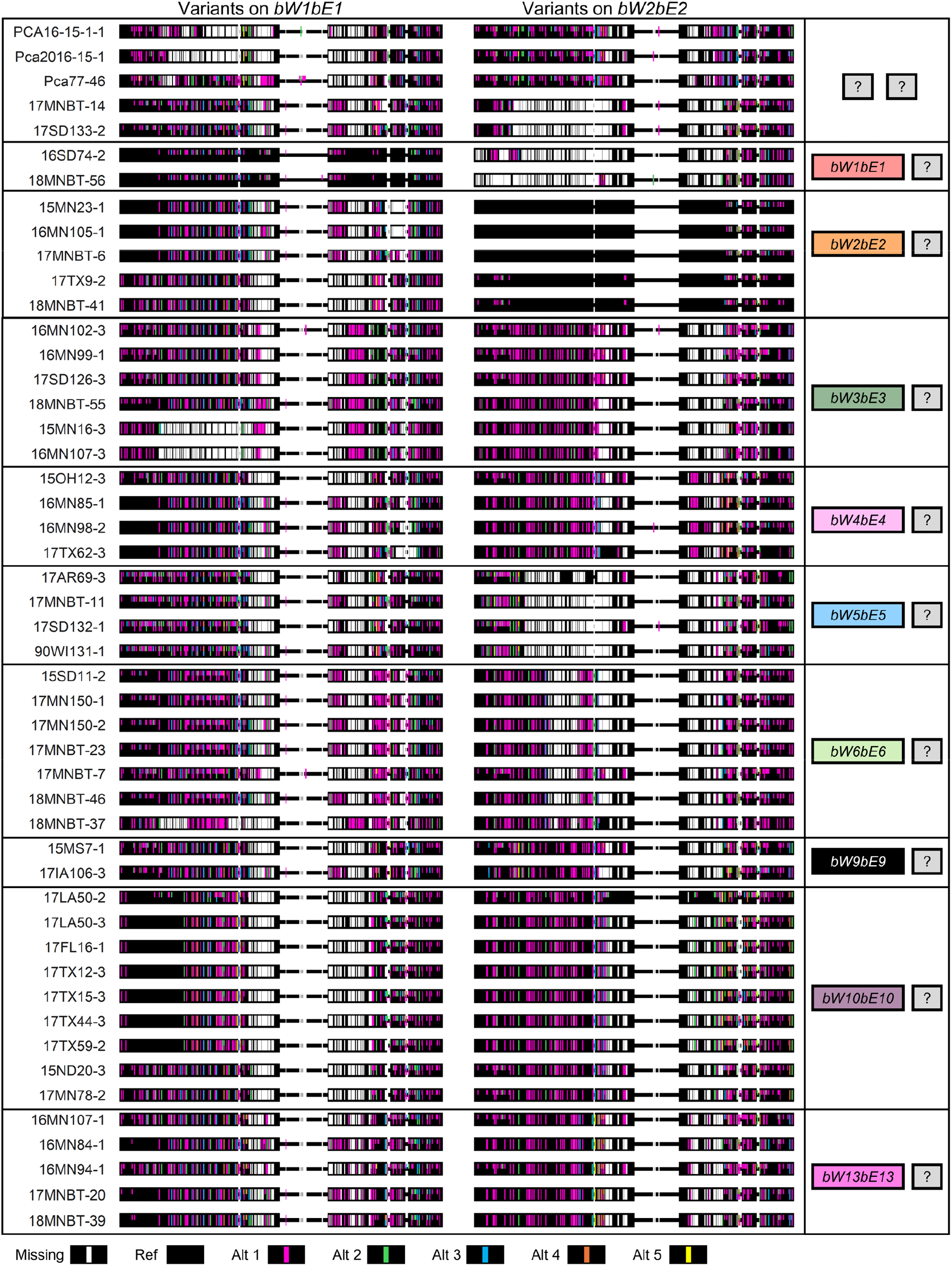
Variants called for 49 *Puccinia coronata* f. sp. *avenae* isolates with one or two unknown *HD* locus alleles in the *HD* locus regions of Pca203 hap1 (*bW1bE1*) and hap2 (*bW2bE2*). Line colors indicate missing (white), reference (black) and alternative (pink, green, blue, orange, yellow) genotypes. Half-length lines indicate heterozygous sites and full-length lines indicate homozygous sites.

**Figure S8.**
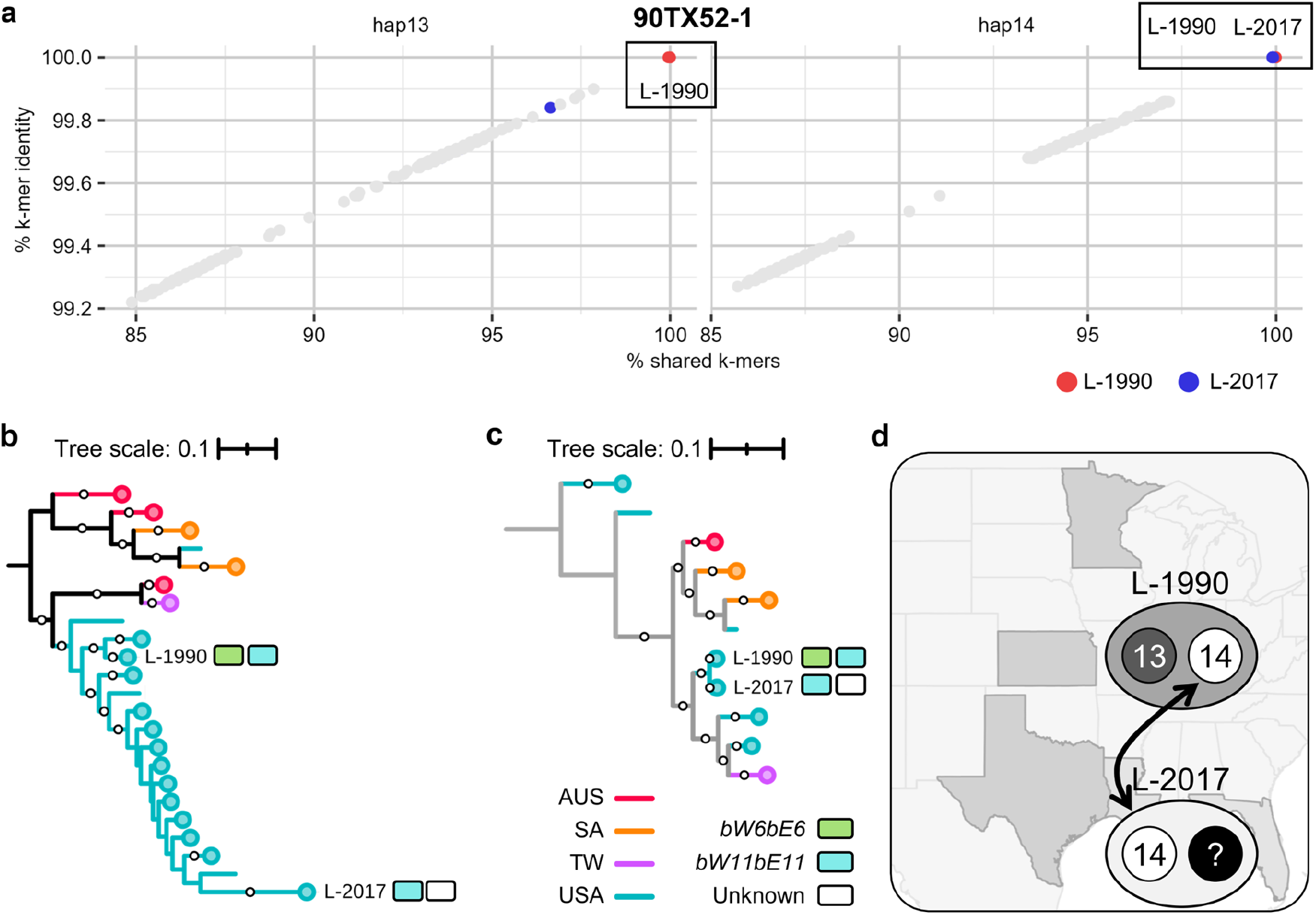
**a)** Plot of % *k*-mer identity (y-axis) versus % shared *k*-mers (x-axis) for 352 *Puccinia coronata* f. sp. *avenae* isolates on hap13 and hap14. Colors indicate relevant lineages; light grey points are all other isolates. **b-c)** Maximum likelihood phylogenetic trees constructed from variants from individual haplotypes **b**) hap13 (194,540 SNPs), **c)** hap14 (209,542 SNPs) with percentages for 500 bootstraps shown on branches. Tree branches are colored by country of origin: AUS = Australia; SA = South Africa; TW = Taiwan; USA = United States of America. Collapsed clades are indicated by circles at tips. *HD* alleles are shown as colored rectangles next to relevant clonal lineages. Tree scales are mean substitutions per site. **d)** Diagram of the proposed relationship between USA lineages and haplotypes involved in somatic hybridization.

**Figure S9.**
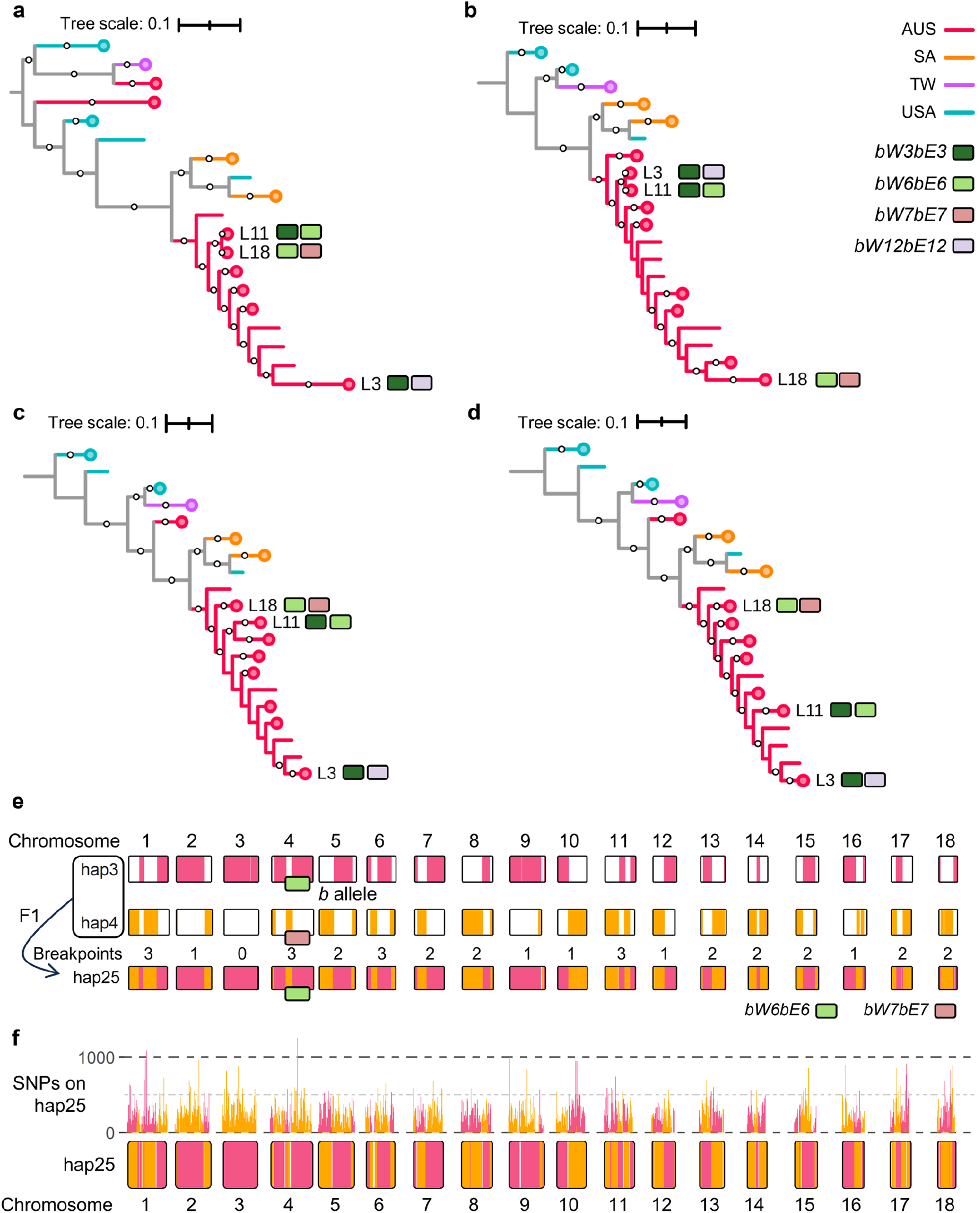
Maximum likelihood phylogenetic trees for 352 *Puccinia coronata* f. sp. *avenae* isolates constructed from variants from individual haplotypes **a**) hap25 (205,993 SNPs) **b**) hap26 (188,803 SNPs) **c**) hap3 (228,300 SNPs) **d**) hap4 (226,617 SNPs). Bootstrap supports 80% or higher are shown as circles at branch midpoints. Tree branches are colored by country of origin: AUS = Australia; SA = South Africa; TW = Taiwan; USA = United States of America. Collapsed branches are indicated by circles at tips. *HD* alleles are shown as colored rectangles next to relevant clonal lineages. Tree scales are mean substitutions per site. **e)** Genome alignment between 20NSW19 haplotypes (hap3, hap4) and hap25 from 20WA94. Chromosomes for the three haplotypes are shown with high-identity alignments shown with colored fill (hap3 = pink, hap4 = orange). **f)** Non-reference variant counts within 100 kb bins from hap3 (pink) and hap4 (orange) on hap25 are shown in the histogram above hap25 chromosomes. Hap25 chromosome fill color is determined by identifying bins with a low density (<50 SNPs/100 kb) of non-reference variants from hap3 (pink) and hap4 (orange).

**Figure S10.**
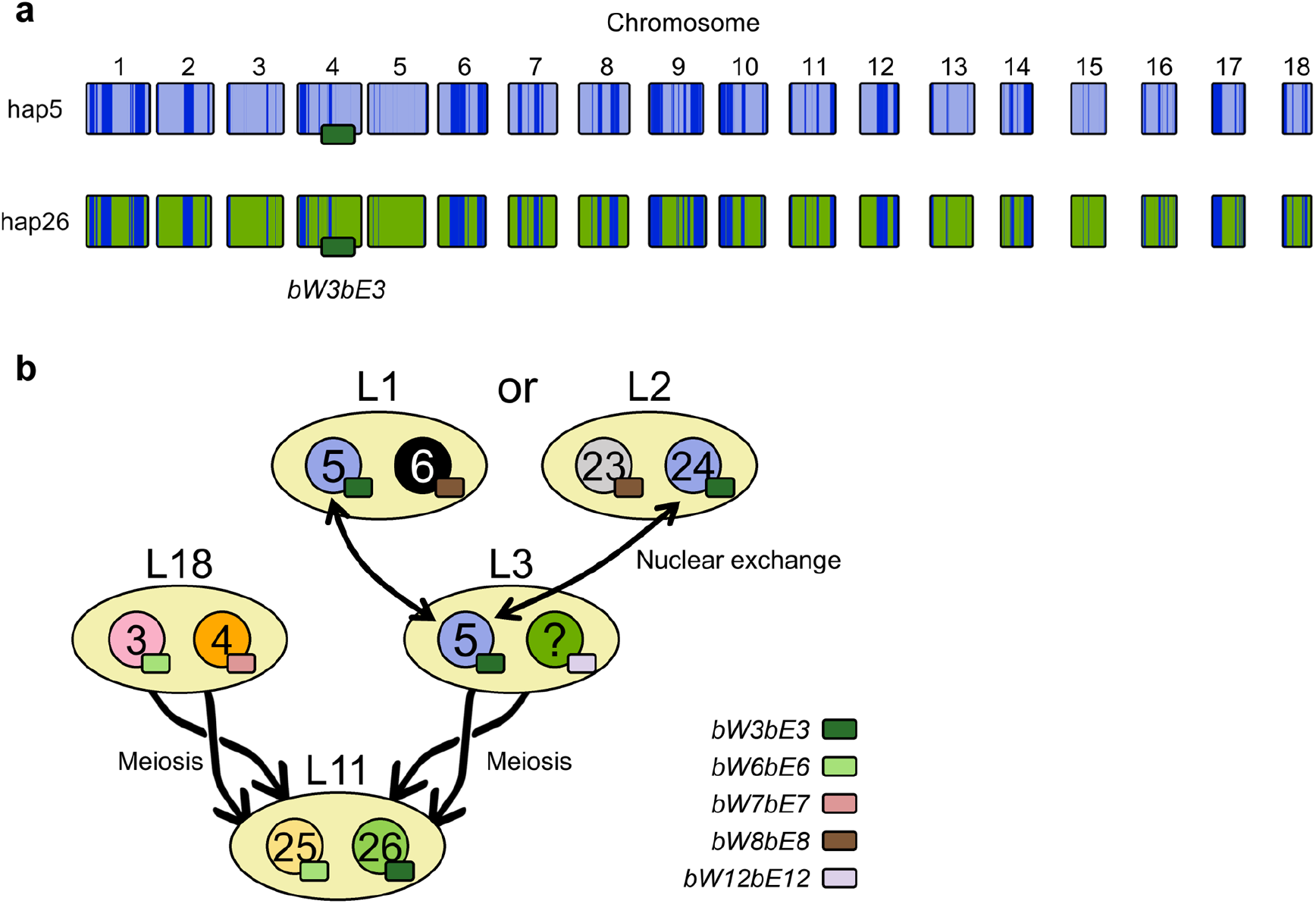
**a)** Regions of 20QLD86 hap5 against 20WA94 hap26 with over 95% identity when aligned are shown in dark blue. Unique sequences are shown in light blue (hap5) and green (hap26). **b)** Considering the mating type composition and shared sequences between hap5 and hap26, it is possible that hap26 from L11 is the progeny of hap5 and an unknown haplotype from L3. L3 may have been founded by somatic hybridization with L1 or L2.

**Figure S11.**
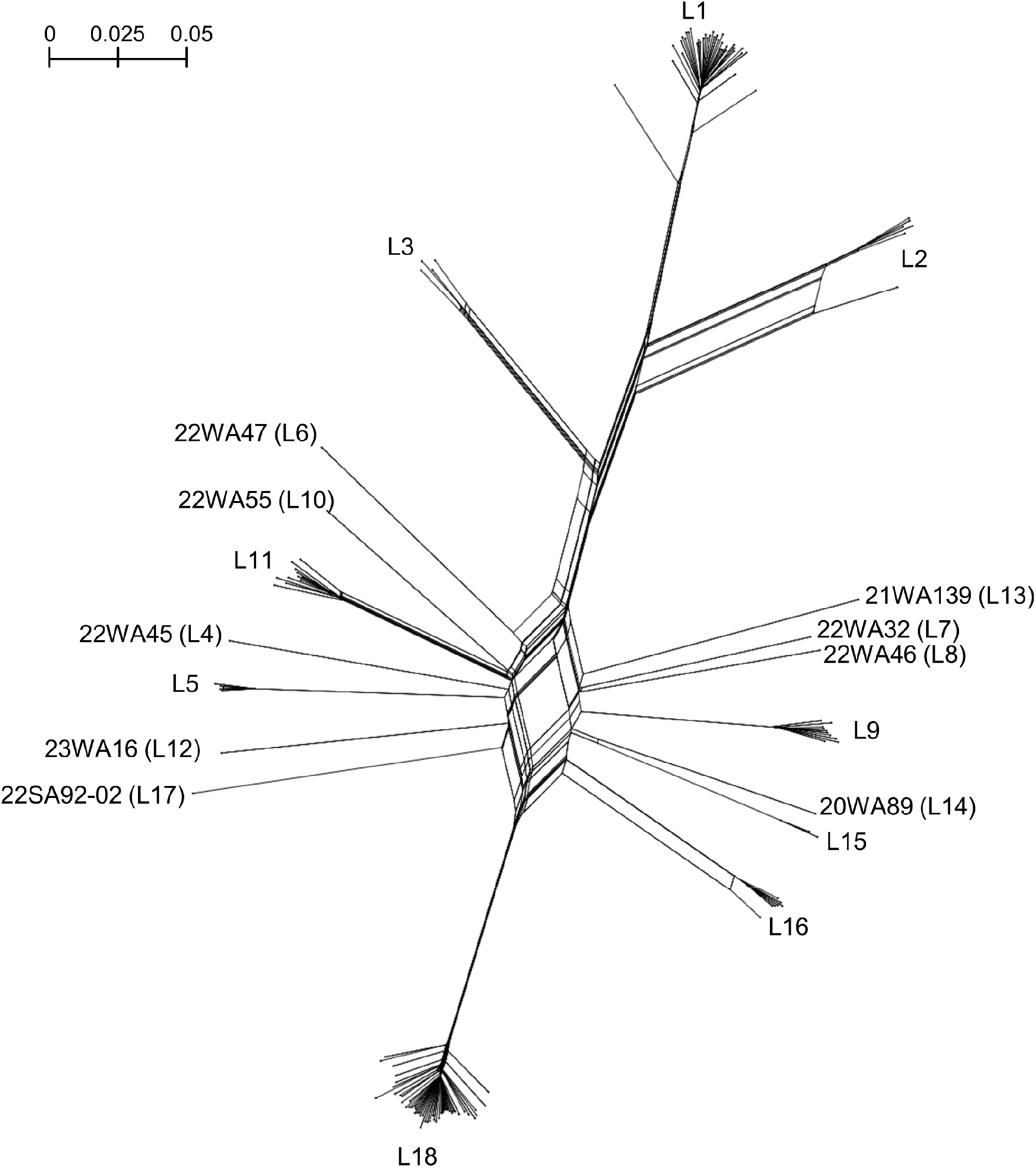
Splitstree network of 137 Australian *Puccinia coronata* f. sp. *avenae* isolates from 18 lineages generated from 391,118 SNPs from the entire Pca203 genome. Heterozygous sites were converted to missing. Scale indicates nucleotide substitutions per site.

**Figure S12.**
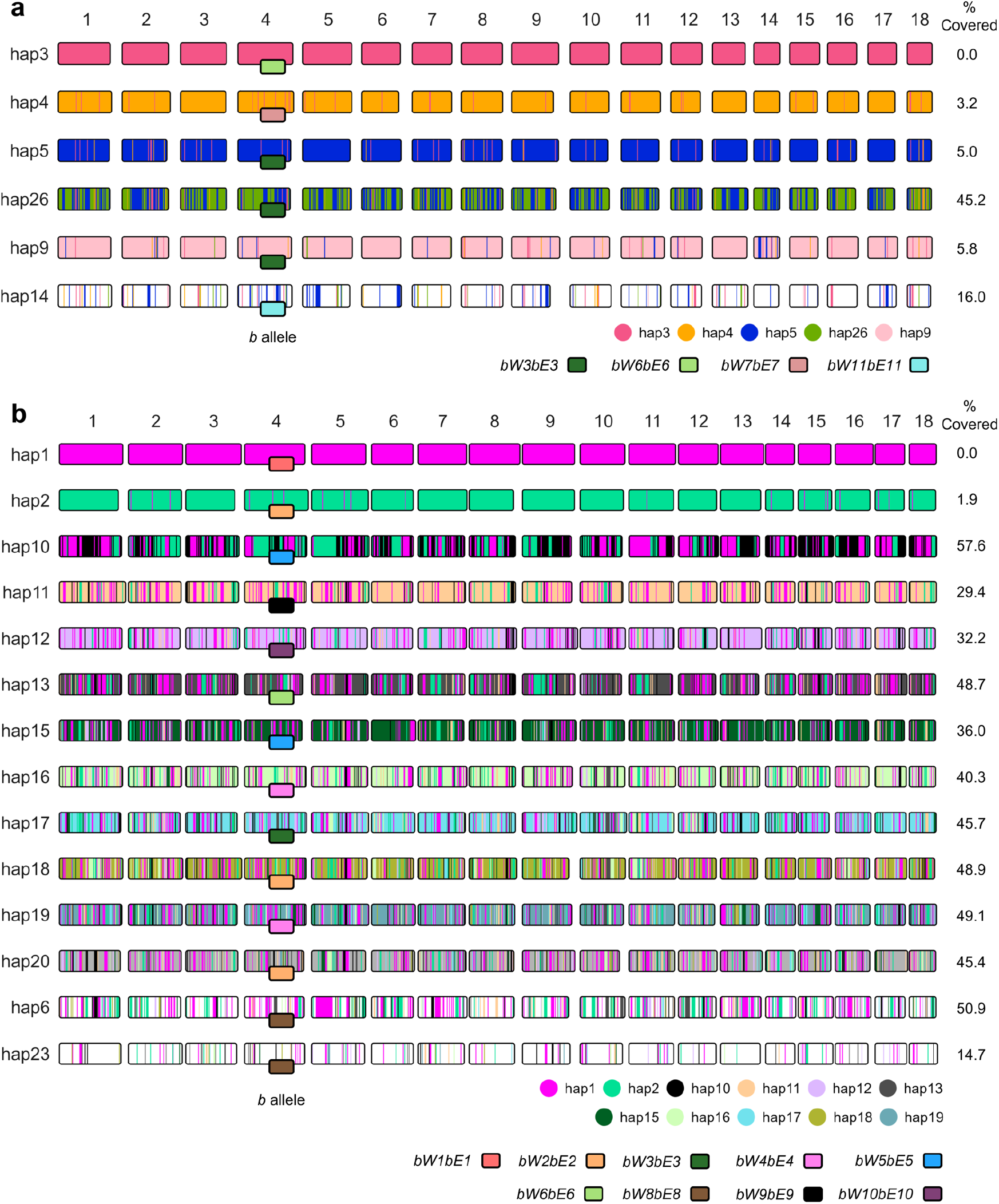
**a)** Shared haplotype blocks across six *Puccinia coronata* f. sp. *avenae* (*Pca*) haplotypes from 100 kb bins with less than 50 non-reference variants which were assigned hierarchically across hap3, hap4, hap5, hap26, hap9, and hap14. Shared sequences with hap3 were assigned across the other five haplotypes first, after which the unassigned areas of hap4 were evaluated on the four haplotypes below in the hierarchy, and so on. Regions in hap14 were not assigned to other haplotypes, so the unassigned hap14 sequence has a white fill. **b)** Generated the same way as panel **a)**, except shared haplotype blocks are assigned across 14 *Pca* haplotypes (hap1, hap2, hap10, hap11, hap12, hap13, hap15, hap16, hap17, hap18, hap19, hap20, hap6, hap23) with hap1 being the first in the hierarchy. Regions from hap6 were not assigned across hap23, so both haplotypes have unassigned sequences filled in white. Chromosome fill color represents unassigned regions or regions shared with haplotypes earlier in the hierarchy. Percent coverage by preceding haplotypes is shown on the right side. *HD* locus alleles indicated by fill color of rectangles at the chromosome 4 midpoint.

## Notes

https://data.csiro.au/collection/csiro:61932

https://github.com/henni164/Pca_pangenome

### Competing Interest Statement

The authors have declared no competing interest.

### Summary of Updates

Acknowledgement of rust sample submitters in the acknowledgements section

