## Supplementary material for "A high-resolution haplotype pangenome uncovers somatic hybridization, recombination and intercontinental migration in oat crown rust": Methods S

Methods S1

Susceptible oat cultivars that were used for amplification vary across countries due to differences in environmental conditions that affect host susceptibility. ‘Marvelous’ was used in the USA, ‘Swan’ in Australia, and ‘NTU Selection 1’ and ‘Swan’ were used in Taiwan. For revival of field samples, the submitted infected leaves were cut into fragments and combined with 1-2 mL Novec oil. The oil mixture was pipetted onto susceptible oat genotypes and the plants were air dried for 30 minutes, misted with water, and kept in humidity chambers (90-99% humidity) for two days before removal to growth cabinets at 23C° for 16 hours light and 18C° for 8 hours dark. After nine days, rust was collected. To isolate single pustules from bulked samples, plants were treated with 15 mL maleic hydrazide per pot and a dilution of 0.5 mg spores in 600 uL of Novec oil was pipetted onto susceptible plants at 9 days of growth. Misting and humidity chamber treatments were the same. Plants were trimmed at around seven days post inoculation (dpi) to keep only leaves with sparse single pustules. Wax paper was scraped against the pustule and leaf surface for collection; these isolations were immediately used to infect new plants. Subsequent infections were performed with approximately 20 mg spores in 400 uL of Novec oil with the same conditions as described before.

For phenotyping assays, 5-6 seeds per genotype were planted with four genotypes per pot. At 9 days post inoculated the plants were treated with 15 mL maleic hydrazide per pot and misted with water, and then a mixture of 50 mg spores with 150 mg talcum powder was spread onto the plants through Miracloth (Millipore Calbiochem®). At 10-11 dpi, infection types were recorded. Multiple infection types may be observed on a single leaf or across biological replicates of the same genotype. As long as infection types do not segregate across the resistant/susceptible boundary, the most common infection type is taken. For example, if four of five leaves have primarily fleck (;) infection types and one leaf has minor sporulation (1), the infection type recorded is fleck. If infection types segregated between resistance and susceptibility across biological replicates (i.e. three plants with score of ‘4’, two plants with score of ‘1’), we repeated the phenotyping for that genotype to ensure an accurate score.

Methods S2

Spores for Hi-C were prepared by suspending 150-200 mg of urediniospores in 13 mL 1% formaldehyde, which was incubated for 20 minutes with periodic vortexing. Glycine powder was added to 1g/100 mL concentrated to quench crosslinking with a 15 minute incubation, again with periodic vortexing. The tubes were spun at 1000 g for two minutes and as much liquid as possible was removed taking care to avoid the clumps of rust spores, which are hydrophobic and tend to float to the top of solutions. The spores were rinsed with 10 mL of water and spun as before. As much water as possible was removed by pipetting. Treated spores were ground with a chilled mortar and pestle with liquid nitrogen added added periodically to maintain cold temperature. Samples were kept at -80 and shipped on dry ice to library preparation providers.

Hi-C libraries for isolates 18MNBT34, 18MNBT36, 18MNBT50, 90AR100, 90MN4b, 90TX52, 20NSW19, 20QLD86, and 20WA72 were prepared at Phase Genomics, Seattle WA USA and sequenced with Illumina Novaseq by Azenta Life Sciences (formerly Genewiz). Libraries for 20WA94, 21ACT116, and 21WA134 were prepared and sequenced with Illumina Novaseq at the Ramaciotti Centre for Genomics, NSW Australia. Hi-C for 20WA95 was not prepared, as it is a clone of 20WA72. During the genome assembly pipeline, Hi-C data from 20WA72 was used with 20WA95 HiFi data.

Methods S3

Initial genome assemblies of 20WA89, 21ACT116, and 21WA134 were larger than expected and showed more fragmentation than assemblies for other isolates performed with the same procedures. To identify potential contaminant isolates, extra contigs (i.e. third copy of segments from main chromosome sequences) were screened with mash v2.0 (Ondov *et al*., 2019) against short-read data for individuals from different lineages. Those with the highest containment across the greatest number of extra contigs were chosen for use in filtering HiFi reads.

Each HiFi read was sketched individually (-s 1000) with mash (v2.0) screen and Illumina reads for the suspected contaminants were screened against the HiFi reads (Ondov *et al*., 2019). *k*-mer identity cutoffs were then applied and tested iteratively by creating a draft assembly with the cleaned reads to assess the success of removal, and the parameters resulting in the best test assembly were chosen to generate the filtered read set. Filtering was deemed adequate when diploid assembly size and contiguity with the filtered reads was comparable to that of pure samples. Percent shared *k*-mers were not considered in filtering as the sequences represented by HiFi and short read sequencing are not expected to overlap perfectly.

Final HiFi read filtering parameters were as follows:

- 20WA89 HiFi reads were screened with 20WA89, 22WA47, and 22WA32 Illumina data and were retained if 20WA89 *k*-mer identity was highest or > 0.9998.
- 21ACT116 HiFi reads were screened with 21ACT116, 21WA134, and 20WA94 Illumina data and were retained if 21ACT116 *k*-mer identity was highest or > 0.9999.
- 21WA134 HiFi reads were screened with 21WA134, 22WA15, and 20WA72 and were retained if 21WA134 or 22WA15 *k*-mer identity was highest or > 0.9999.

Approximately 21% of 20WA89 HiFi reads longer than 10 kb were removed with the remaining reads randomly downsampled to 40X coverage. For 21WA134, 11% of the reads longer than 10 kb were removed and remaining reads were downsampled to 30X coverage. Finally, 38% of 21ACT116 reads longer than 10 kb were removed and downsampled to around 30X coverage. Downsampling was performed with seqkit v2.7.0 (Shen *et al*., 2016).

Methods S4

PacBio HiFi reads for all isolates were input into hifiasm v0.16.1 (Cheng *et al*., 2021) with their respective Hi-C reads integrated for all samples except 20WA95, for which the Hi-C data from its clonal relative 20WA72 was used. For 20WA89, 20WA94, 21ACT116, 21WA134, and 21WA139, the same was performed with hifiasm v0.19.5 (Cheng *et al*., 2021). Haplotypes resulting from the hifiasm assembly for each isolate were combined and contigs belonging to the mitochondrial genome were filtered using BLAST+ v2.13.0 (Camacho *et al*., 2009). The PacBio reads were mapped back to the assembly using minimap2 v2.22 (--ax --secondary=off) (Li, 2018) and contig coverage was calculated with bbmap (v39.01) pileup (sourceforge.net/projects/bbmap/). Contigs with average coverage of <= 5X were removed, except 20WA89, 20WA94, 21ACT116, and 21WA134, which used a cutoff of <= 10X. The remaining contigs were then BLASTed to the NCBI nucleotide database with blast+ v2.13.0 and contigs without ‘*Puccinia*’, ‘*Medioppia*’, ‘*Phakopsora*’, ‘*Melampsora*’, ‘*Uromyces*’, or ‘ribosomal’ in their top five hits were removed (Camacho *et al*., 2009; Sayers *et al*., 2022). The cleaned assembly was prepared and phased with NuclearPhaser v1.1 (<https://github.com/JanaSperschneider/NuclearPhaser>). Scaffolding was completed by aligning each haplotype to the Pca203 ‘A’ genome with D-Genies v1.5.0 to determine contig order and orientation, in addition to generating Hi-C contact maps with HiC-Pro v3.1.0 and hicexplorer v3.7.2 (Servant *et al*., 2015; Cabanettes & Klopp, 2018; Ramírez *et al*., 2018). Unplaced contigs were excluded from the analysis of haplotypes as they were already represented in the chromosomes, were on average short (8,024- 633,353 bp, mean = 43.83 kb) and had high repeat content (44.77-85.35%, mean = 68.30%) (Figure S1; Table S4).

Repeats in the scaffolded genomes were masked with repeatmodeler v2.0.2a and repeatmasker v4.1.2pl (--nolow), retaining only classified repeats (Smit *et al*., 2015; Flynn *et al*., 2020). RNAseq reads from 12NC29 (spores, haustoria) and Pca203 (5 dpi) were used (Miller *et al*., 2018; Henningsen *et al*., 2022). RNAseq reads from 12NC29 and *Pca*203 were mapped to individual haplotypes and unplaced contigs separately with Hisat2 v2.2.1 (--max-intronlen 3000 --dta --no-unal --rna-strandedness RF) (Miller *et al*., 2018; Kim *et al*., 2019; Henningsen *et al*., 2022). Read mappings were merged and used as input for Trinity v2.13.2 in genome-guided mode (--jaccard_clip --genome_guided_max_intron 3000 --SS_lib_type RF) (Grabherr *et al*., 2011). Hisat2 v2.2.1 (--max-intronlen 3000 --dta --no-una) was also used to align the RNAseq reads in preparation for assembly with stringtie v2.2.1 (-s 1 -m 150) (Pertea *et al*., 2015; Kim *et al*., 2019). Codingquarry v2.0 was run on infection and spore transcripts separately (Testa *et al*., 2015) and filtered as described in Sperschneider et al. (2023). Funannotate v1.8.5 training was run on previous Trinity transcripts (--stranded RF --no_trimmomatic --jaccard_clip) (Palmer & Stajich, 2020). Funannotate v1.8.5 predict was run using on the deduplicated and repeatmasked genomes with the Trinity, coding quarry, and pucciniomycotina EST evidence (--ploidy 2 --optimize_augustus --busco_seed_species ustilago --weights pasa:10 codingquarry:0). Funnanotate v1.8.13 update was run with this training information. Finally, annotations were finalized by processing with transdecoder v5.5.0 and agat v1.0.0 (Haas, BJ <https://github.com/TransDecoder/TransDecoder>; Dainat, 2023).

Pca203-*STE3.2.2* was not annotated correctly due to an overlapping gene annotated on the opposite strand; the correct gene model for *STE3.2.2* was recovered from augustus predictions and confirmed by alignment to *Puccinia graminis* f. sp. *avenae* alleles with CLUSTALW (Larkin *et al*., 2007). *STE3.2.2* required recovery and/or correction in nearly all isolates and some haplotypes had incorrect exon boundaries in *STE3.2.3* which were corrected. The 5’ ends of *bW-HD1* and *bE-HD2* are highly variable and because RNAseq data from 12NC29 and Pca203 were used for annotation of all haplotypes, several gene models were recovered from augustus, coding quarry, or pasa predictions and/or manually corrected using conserved domain information and protein alignments. Preliminary screens indicated that one *HD* allele pair for the published reference 12NC29 (Miller *et al*., 2018) was not sampled in any of the chromosome-level references; as such, the missing allele was identified in the 12NC29 contig-level reference genome, and the locus was added to the screening.

Methods S5

The cactus-pangenome pipeline generates a pangenome graph and can also call variants with each input genome as the “reference”, using the other genomes as “samples”. By listing every haplotype as a “reference”, 32 VCF files were generated that map variants to physical positions along each haplotype. Variants along each reference were binned every 100 kb and bins with <= 50 non-reference SNPs were marked as “shared” between the reference and the sample haplotype. This threshold was determined by visualizing non-reference SNPs distributions across hap25, where hap3 and hap4 are samples and hap25 is the reference. Histogram bins without any SNP calls were ignored to prevent repetitive or poorly aligned regions from being assigned as shared.

Adjacent bins that were marked as shared for each sample-reference pair were combined with bedtools (v2.31.1) merge (Quinlan & Hall, 2010). The intermediate files produced from this are pairwise shared haplotype blocks between all haplotypes. However, to produce a sensible visualization, a hierarchy must be applied to handle overlaps between shared haplotype blocks. For example, hypothetical haplotypes hapA and hapB have a region in common. HapC also shartes part of this region and shares a different region with hapB. In a hierarchy, we could place hapA at the top, so the sequence shared by all three haplotypes is colored as “hapA” and only the non-overlapping section of hapC is colored as “hapB” (Figure 1b). The order of the hierarchy we used was determined by calculating the proportion of each haplotype covered by others without removing overlaps, which is shown in Table S9. Using a custom script, these regions were then subjected to an ordered bedtools subtraction, whereby regions from the first haplotype in the hierarchy are assigned first across all other haplotypes. These regions are added to the consensus file that is subtracted against for the second haplotype in the hierarchy, and so on (Quinlan & Hall, 2010; <https://github.com/henni164/Pca_pangenome/tree/main/Figure5>).

After all regions are assigned hierarchically, those <= 50 Kb were filtered out and the remaining regions were visualized with R package ggplot2 (Wickham, 2016).

For the analysis of Australian haplotypes, parent haplotypes hap3 and hap4 were chosen as the first two haplotypes for hierarchical recombination block assignment. Hap5 was third for being divergent from hap3 and hap4, and its recombinant hap26 was placed in fourth position. Finally, hap6 and hap23, were placed in fifth and sixth position to highlight their divergence from other Australian haplotypes. Regions with low non-reference SNP density from these first six haplotypes were assigned across the other Australian haplotypes (hap25, hap7, hap8, hap27, hap28, hap29, hap30, hap31, hap32).

For the analysis of US haplotypes, haplotypes from Pca203 were used first as it is likely the oldest isolate in the collection and caused significant epidemics in the 1940s (Stoa & Swallers, 1950), hap1 and hap2 regions were assigned first across USA haplotypes, hap6, and hap23. Shared sequences from the other USA haplotypes were assigned in descending numeric order with hap6 and hap23 in the final positions. Hap9 and hap14 were not included in this set due to their divergence to the other US haplotypes. Instead, they were visualized with hap3, hap4, hap5, and hap26 regions, as they share more sequence with Australian haplotypes.
